# Factors Shaping Young and Mature Bacterial Biofilm Communities in Two Drinking Water Distribution Networks

**DOI:** 10.1101/2021.03.10.434709

**Authors:** Dan Cheng, Mats Leifels, Carlo Miccolis, Stefan Wuertz, Janelle R. Thompson, Ulrich Szewzyk, Andrew J. Whittle

**Author notes:** Corresponding author Singapore Centre for Environmental Life Sciences Engineering (SCELSE), Nanyang Technological University, 60 Nanyang Drive, SBS-01N-27 Singapore 637551.

## Abstract

The presence of biofilms in drinking water distribution systems (DWDS) can affect both water quality and system integrity; yet these systems remain poorly studied due to lack of accessibility. We established two independent full-scale DWDS Testbeds (A and B) on two different campuses situated in a tropical urban environment and equipped them with online sensors. Testbed B experienced higher levels of monochloramine and lower water age than Testbed A within the campus. Based on long amplicon-sequencing of bacterial 16S rRNA genes extracted from the mature biofilms (MPB) growing on pipes and young biofilms (YSB) growing on the sensors, a core community was identified in the two testbeds. The relative abundances of operational taxonomic units at the family level, including *Mycobacteriaceae, Methylobacteriaceae, Rhodospirillaceae, Nitrosomonadaceae,* and *Moraxellaceae,* were consistent for MPB and YSB on each campus. The MPB community was found to be influenced by conductivity, sample age, and pipe diameter as determined by both canonical correlation analysis and fuzzy set ordination. MPB displayed higher α-diversity based on Hill numbers than YSB; in general, second order Hill numbers correlated positively with conductivity and sample age, but negatively with ORP and nitrite. *Pseudomonas* spp. together with *Bacillus* spp. likely initiated biofilm formation of YSB on Testbed A under conditions of reduced monochloramine and high water age. Significant levels of orthophosphate were detected in YSB samples at two stations and associated with higher levels of stagnation based on long-term differential turbidity measurement (DTM). Orthophosphate and DTM may act as indicators of the biofilm growth potential within DWDS.

**Highlights:** - Established two testbeds to study biofilms in full-scale distribution system
- Biofilms on pipes and sensors had core community
- Temporal effect and higher α-diversity for biofilms on pipes
- Water chemistry was related to biofilm community differences

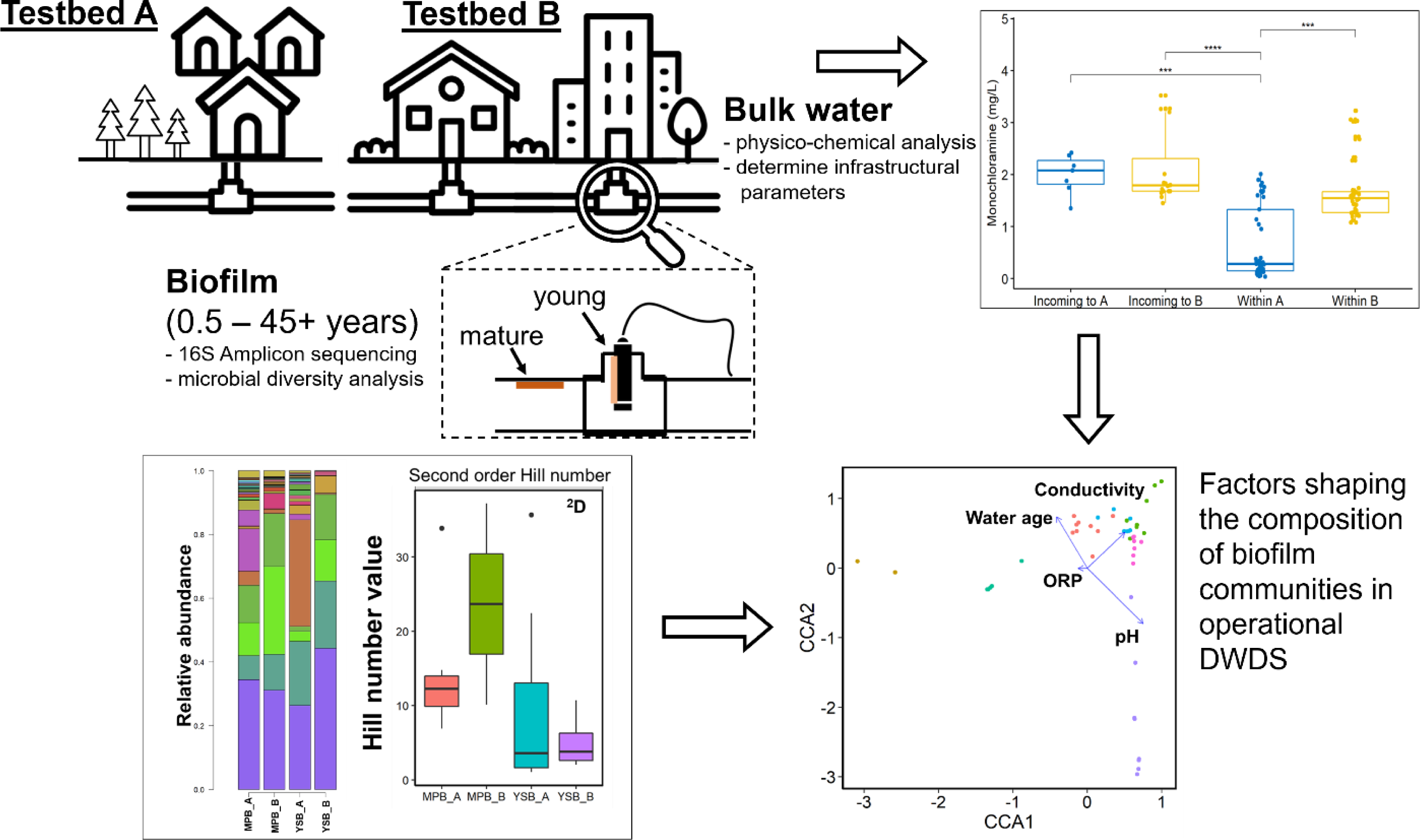
Graphical Abstract

## 1. Introduction

There is an increasing recognition of public health risks in drinking water distribution (DWDS) and premise plumbing systems where water stagnation and decay of residual disinfectant favor regrowth of microorganisms within the pipe networks (Allaire et al. 2018, Hull et al. 2019, Raskin and Nielsen 2019). Biofilms can act as reservoirs of both enteric and opportunistic pathogens, which may then be released back into the bulk water (Cruz et al. 2020, Flemming et al. 2016, Flemming and Wuertz 2019). Several outbreaks of illnesses such as legionellosis, which results in high morbidity and mortality in susceptible individuals, have been associated with poor drinking water quality (Ashbolt 2015). A prime example is a recent outbreak of Legionnaire’s disease (a severe manifestation of legionellosis) in the City of Flint, Michigan, which was attributed to the release of bioavailable iron into the water stream after switching to a more corrosive water source (Schwake et al. 2016).

While cultivation-based methods are considered the gold standard for examining microbial drinking water quality, various molecular approaches have been proposed to account for viable but non-cultivable (VBNC) bacteria that have been described to reside in DWDS biofilms (Dwidjosiswojo et al. 2011, Konigs et al. 2015). More than 98% of the total biomass present in DWDS have been found to be present as biofilms and loose deposits in an unchlorinated DWDS as quantified by adenosine triphosphate (ATP) analysis and DNA-pyrosequencing (Liu et al. 2014). There is limited access to operating water main pipes and hence, most prior studies have been conducted on premise plumbing systems including i) mature pipe biofilms in buildings designated to be demolished (Henne et al. 2012), ii) biofilms within replaced water meters (Ling et al. 2016), and iii) young sentinel biofilms growing on carrier coupons in model systems (Douterelo et al. 2014) or fire hydrant outlets (Douterelo et al. 2020).

While offering unique insights, such studies have shown that biofilms forming under various conditions and on different substrate materials can host significantly different microbial communities and are therefore not fully representative of operational large-scale DWDS. For example, Buse et al. (2014) found that the families *Burkholderiaceae*, *Characeae*, *Epistylidae*, *Goniomonadaceae*, *Paramoebidae*, *Plasmodiophoridae*, *Plectidae*, *Sphenomonadidae*, and *Toxariaceae* were predominant in biofilms on unplasticized Polyvinyl Chloride (uPVC) substrates and *Enterobacteriaceae*, *Erythrobacteraceae*, *Methylophilaceae*, *Acanthamoebidae*, and *Chlamydomonadaceae* in biofilms on copper pipes. While offering insights into the dynamics of DWDS biofilms, most of the novel testbed systems referred to pilot-scale plants with re-circulating water. Consequently, there are few published data on biofilm composition during early biofilm development within an operational water distribution network.

The availability of nutrients such as carbon, nitrogen and phosphorus has been demonstrated to be essential for high metabolic potential affecting both rates of biofilm growth and microbial community composition. Kasahara et al. (2004) showed that dechlorinated tap water amended with 50 µg/L of phosphate (in KH_2_PO_4_ to increase the content of orthophosphate, hereafter referred to as PO_4_-P) could increase biofilm growth on a pipe wall in a continuously rotating model reactor. Similarly, Fang et al. (2009) found the addition of 30 and 300 µg/L of PO_4_-P could induce thicker and less homogeneous biofilms with more biomass in a laboratory reactor. However, little is known about the effect of biofilm development on the nutrient concentration in the bulk water of an operational DWDS system.

Although online monitoring of biofilms has been proposed for DWDS since 2000 (Flemming et al. 1998, Flemming et al. 2011, Janknecht and Melo 2003, Strathmann et al. 2013), water utilities have been slow to respond due to the high costs of installation and maintenance (Allaire et al. 2018, Carminati et al. 2020). Traditionally, biofilm formation has been associated with the detection of a deposit on a surface using differential turbidity measurement (DTM) between the clean and colonized probe (Flemming et al. 1998). The online WaterWise system was initially developed to detect leakages but has also enabled additional real-time monitoring of water quality parameters, including pH, ORP, temperature, conductivity and turbidity, for the whole DWDS in Singapore (Allen et al. 2013, Kitajima et al. 2020, PUB 2016). Kitajima et al (2020) suggested that water age and monochloramine drive the community of both biofilm and bulk water samples. Here we report on the use of these sensors to observe the microbial community dynamics in biofilms from two operational campus drinking water networks, referred to as Testbeds A and B. Both Testbeds are supplied with piped water from the national drinking water distribution system that has been treated with similar levels of monochloramine as secondary disinfectant. The testbeds differ in how the water is stored and distributed. A high-resolution shot-gun sequencing analysis of bacterial 16S rRNA genes was used to analyze the community structure associated with biofilms of different ages and from different environments: i) mature pipe biofilms (MPB) collected from coupons (interior surfaces of old/extant pipes) and ii) young sensor biofilms (YSB) that developed on WaterWise water quality sensors installed inside the pipes at specific locations within each Testbed. We explored temporal and spatial effects on the communities of YSB and MPB and interpreted sequencing data using operational conditions and physicochemical water quality parameters.

## 2. Materials and methods

### 2.1 DWDS Testbeds

We established two campus testbeds that receive piped drinking water (bulk water) from the national supply grid with monochloramine as residual disinfectant. Each campus has a resident population of about 40,000 people with an average daily water demand of about 6,000 m^3^·d^-1^ and 4,600 m^3^·d^-1^ for Campus A and B (in 2016), respectively (Figure S1A). Pipes varied in age from 14 to 47 years and were manufactured from ductile iron either unlined or lined with cement or polyurethane (Table 1).

**Table 1.** Characteristics of Campus DWDS

| Parameter (unit) | Campus A | Campus B |
| --- | --- | --- |
| Fire hydrant system | Tapped from domestic line | Separate loop |
| Pipe material | DI<sup>1</sup> with or without cement lining | DI<sup>1</sup> with cement or polyurethane lining |
| Pipe diameter (mm) | 300 and 230 | 250 and 200 |
| Pipe (biofilm) age when sampled (y) | 34 - 47 | 21 & 14 |
| Temperature (°C) | 29.7 ± 0.3 | 31.3 ± 0.5 |
| Daily flow consumption (TCMD<sup>2</sup>) | 0-3.7 | 0-3.0 |
| Network pressure (m) | 30 - 65 | 10 - 40 |
| Inlet<sup>3</sup>: Monochloramine (mg/l) | 2.00 ± 0.38 | 2.12 ± 0.72 |
| Within campus: Monochloramine (mg/L) | 0.66 ± 0.58 | 1.70 ± 0.59 |
| ORP (mV) | 200 - 460 | 336 - 460 |
| DTM (FNU) | 0.14-2.01 | 0.01-1.05\| |
| Relative water age (h)<sup>4</sup> | 80 - 164 | 20 – 30 |
| Total particle count (per ml) | (3.40 ± 1.22) × 10<sup>5</sup> | (2.27 ± 1.64) × 10<sup>5</sup> |
| Total bacterial count (per ml<sup>5</sup>) | (8.75 ± 2.51) × 10<sup>4</sup> | (5.93 ± 3.75) × 10<sup>3</sup> |

<sup>1</sup>DI – ductile iron
<sup>2</sup> TCMD – thousand cubic meter per day
<sup>3</sup> Stations A11 and B13/B14 are the inlets receiving city water for campus A and B, respectively.
<sup>4</sup> Defined by hydraulic model relative to inlet conditions.
<sup>5</sup> Measured by flow cytometry with live/dead staining kit.

Each testbed uses a network of fixed WaterWise sensor nodes, equipped for continuous monitoring of hydraulic (flowrate, fluid pressure and acoustics) and basic environmental variables relating to water quality (temperature, conductivity, pH and ORP, and turbidity) using a YSI EXO2 sonde (multi-parameter probe) manufactured by Xylem (Rye Brook, NY, USA) (Allen et al. 2013).

Table 1 summarizes the characteristics of the two campus DWDS together with the averaged water quality indicator parameters. The measured concentrations of monochloramine in the delivered source water were similar for both campuses (2.00 ± 0.38 mg/L and 2.12 ± 0.72 mg/L for inlet conditions at stations A11and B13/B14, respectively: Figure S1) during February 2017 and July 2018. However, due to the interception and storage of water in a large tank on Campus A (7.5 – 8.0 m deep; Fig. S1a), levels of residual disinfectant within Campus A (0.66 ± 0.58 mg/l) were much lower than on Campus B (1.70 ± 0.59 mg/l). We estimated the relative water age (relative to inlet conditions) on Campus A using a calibrated hydraulic model EPANet (Rossman 2000) to be in the range of 85 - 164 h. In contrast, residual disinfectant levels on Campus B were consistent with a relative shorter water age in the range of 20 - 30 h (Figure S1c). Conductivity levels varied with both source water and upstream treatment. Continuous measurements over a one-year sampling period (August 2017-August 2018) showed some variations within each testbed, but a much lower average conductivity for Campus A (175 ± 26 µS cm^-1^) compared to Campus B (290 ± 32 µS cm^-1^). Fluctuations in conductivity on a weekly or daily basis suggested subtle changes in the mixing of the source water supply.

Prior data from WaterWise sensors installed in stagnant bulk water showed a rapid shift in differential turbidity measurement (DTM) values; it was thus hypothesized that long-term online monitoring of DTM (at its log value) could be an effective indicator of stagnation in the network.

### 2.2 Biomass collection, DNA extraction, and Illumina sequencing

Two types of biofilm were collected at the same locations: 1) mature pipe biofilms (MPB; extracted pipe wall coupons obtained during installation of the sensor nodes) and 2) young sensor biofilms (YSB; sampled during routine sensor maintenance). Figure S1D summarizes the time sequence of sampling operations, mainly phase I (December 2016 - October 2017) and phase II (November 2017 - April 2018) for the MPB samples. YSB samples were obtained from the surfaces of the sensor sondes during regular maintenance or replacement operations (aged from 1- 14 months in all 7 cases).

The pipe coupons were obtained by under-pressure drilling through the pipe wall, with no interruption of water supply. Each pipe coupon was immediately transferred to a sterile petri-dish and brought back to laboratory on ice for processing in less than half an hour. Biofilm samples were collected with sterile cotton swabs by swabbing the inner area of the cut pipe section (surface area = 7,536 mm^2^) after rinsing the surface with sterile phosphorus buffer saline (PBS). Biofilm was removed from the swab by rigorous vortexing for 7 min and then centrifugating at 7,000 g for 15 min at 4°C. The supernatant was removed, and the remaining cell suspension transferred to a pathogen lysis tube (Qiagen, Hilden, Germany) with glass beads for total genome extraction. The sample was mechanically lysed by FastPrep-24 (MP Biomedicals, OH, USA) at a velocity of 6.5 m/s for 10 s with a 2 min intermission for 3 cycles at room temperature. The supernatant was transferred and subjected to RNaseA digestion at 37°C for 30 min before proceeding with the Qiagen UCP pathogen mini kit (Qiagen, Hilden, Germany) according to the manufacturer’s protocol. The extracted genomic DNA was eluted in 35 µL of buffer AVE twice and stored at - 80 °C.

Sequencing was conducted with the Illumina multiplex strategy using an Illumina genome analyzer IIx (Illumina, Singapore) at the Genome Institute of Singapore. Extracted DNA was amplified using the primer pair 338F (5’-ACTYCTACGGRAGGCWGC-3’) and 1061R (5’-CRRCACGAGCTGACGAC-3’) following the protocol developed by Ong et al. (2013). The amplified PCR products (∼700-1,000 base pairs) were purified with the QIAquick PCR purification kit (Qiagen, Hilden, Germany). DNA sequencing libraries were labelled with multiplex indexing barcodes using the Illumina Multiplexing Sample Preparation Oligonucleotide Kit (Illumina, Singapore).

### 2.3 Reconstruction and classification of 16S rRNA amplicon sequences

Full length 16S rRNA gene reconstructions were produced from the short sequencing reads using the *expectation maximization iterative reconstruction of genes from the environment* (EMIRGE) amplicon algorithm (Miller et al. 2011, Miller et al. 2013) and following an updated workflow (https://github.com/CSB5/GERMS_16S_pipeline) described by Ong et al. (2013) in GERMS using R, to determine how the community structure differed between categorical variables. Two-dimensional visualizations of the community structure were conducted using non-metric multidimensional scaling (nMDS) with Bray-Curtis distances at phylum, class, family, genus, and operational taxonomic unit (OTU) levels. Permutational multivariate analysis of variance (PERMANOVA) was applied to assess the similarity/differences in bacterial community structure among samples in the *Adonis* function of the vegan package in R (Anderson 2014-2017). Community differences between samples were calculated as Bray-Curtis dissimilarity and the similarity matrix visualized by principal coordinate analysis (PCoA, also referred to as MDS). Alpha diversity analysis was conducted with R studio and vegan package according to Ling et al. (2016). OTUs were defined by clustering at 3% divergence (97% similarity). Final OTUs were taxonomically classified using BLASTN against a curated database derived from NCBI and Greengenes (all data available upon request).

Computational multivariate analyses were conducted in R according to Roberts (2009) and Torondel et al. (2016). The *bioenv* function of the vegan package was used to determine environmental variables showing a maximum correlation with community dissimilarities and then plotted them as vectors along with the best subset of taxa on the nMDS plot. Canonical correlation analysis (CCA) based on Bray-Curtis distance matrices was used to identify and measure the best associations among two sets of variables that could describe the community structure by using the Adonis function from the vegan library (Torondel et al. 2016). The fuzzy set ordination (FSO) was conducted to test effects of perturbation in environmental variables on the dissimilarity matrix of the biofilm community structure based on the distance index calculated by the Horn method proposed by Boyce and Ellison (2001). Prior to correlation analysis, the OTU table was sub- sampled to the same sequencing depth (2,763 OTUs/sample) for all the MPB and YSB samples. Multiple subplots for all the environmental variables against the second order Hill number (inverse Simpson index) (Chao et al. 2014) were compared in a single plot based on vegan and plyr packages in R. To indicate both the complexity and diversity of the microbial community populating the biofilm, Hill numbers referring to richness index (^0^D), exponential of Shannon diversity (^1^D), and the inverse Simpson index (^2^D) were calculated according to the approach described by Alberdi and Gilbert (2019).

### 2.4 Physico-chemical characterization of bulk water

Concentrations of various nutrients and ions were determined manually in triplicate for the bulk water collected from individual stations on each campus. Chloride and sulphate concentrations were determined by ion chromatography (Prominence HIC-SP, Negeri Sembilan, Malaysia). Concentrations of total organic carbon (TOC), total carbon (TC), total inorganic carbon (TIC), and total nitrogen were obtained by TOC-L analyzer (Negeri Sembilan, Malaysia). Ammonia (NH3-N), nitrite (NO_2_-N), nitrate (NO_3_-N), and orthophosphate (PO_4_-P) were determined with the DR-3900 spectrophotometer (HACH Lange, Düsseldorf, Germany). Chlorine based residual disinfectant concentrations in the water phase were quantified using the HACH DR900 (HACH Lange, Düsseldorf, Germany).

### 2.5 Bacterial activity in water

The activity of hydrolase enzyme unique to bacteria present in bulk water was examined by the BactiQuant (Mycometer, Horsholm. Denmark) water kit according to the manufacturer’s protocol (Reeslev et al. 2011). In brief, fresh bulk water samples of 250 mL were filtered on site through a 0.22 µm filter (Merck-Millipore, MS, USA) and the retentate incubated with substrate for 30 min at room temperature. Analytical blanks were included according to the manufacturer’s instruction as process controls. The BactiQuant Water value (BQW) is obtained from the following dimensional equation:

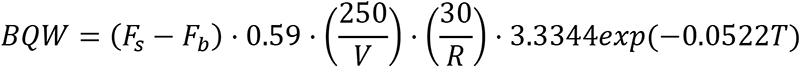

whereas *F_s_* is the sample fluorescence, *F_b_* is the blank fluorescence, V [mL] is the volume of the water sample, R [min] is the reaction time, and T [°C] is the temperature.

### 2.6 Flow cytometry measurement

Flow cytometry (FCM) measurement using the LIVE/DEAD BacLight viability and counting kit (Invitrogen, CA, USA) was used to quantify the live/intact and total bacterial cells in water samples on the Attune NxT flow cytometer (Thermo Fisher Scientific, MS, USA) according to the manufacturer’s protocol. In brief, 1 ml of the sample was stained with 1.5 µL of SYTO9 stain (3.34 mM) and 1.5 µL of propidium iodide (PI) (20 mM) in filtered DMSO and incubated in the dark for 10 min. The FCM analysis was performed in triplicate for each sample at a flow rate of 100 µL min^-1^ and was calibrated to measure the bacterial count of 200 µL of sample. Fluorescence dyes SYTO9 and propidium iodide were excited at a wavelength of 488 nm and emissions measured at 561 nm and 640 nm, respectively.

### 2.7 Statistical analysis

Statistical analysis was conducted in R. The Kruskal-Wallis test was used to assess whether the mean ranks were the same at different stations, in addition to pairwise comparison with Dunn’s Test. Results were plotted using RStudio Version 1.2.5001 / 19 (RStudio, MS, USA) with packages dplyr, ggpubr, rstatix, and outliers.

## 3. Results and Discussion

### 3.1 Comparison of bulk water properties on Testbed A and B

The average live bacterial cell counts in the bulk water at locations in Testbed A were significantly higher than measured in the city water at the inlet (see Table S1). Live cell counts were positively correlated with relative water age and negatively correlated with level of residual disinfectant (monochloramine concentration), rising from an average of 30 cells/mL at the inlet to 60-65 cells/ml at stations A15 and A16, with a concomitant increase in the ratio of live to total cells (1.7% vs 6.3 – 7.4%). A similar trend in enzymatic activity was observed from the BactiQuant Water (BQW) values at these stations (see Table S1). In contrast, there was a lower count of dead cells within the network (A15, A16) compared to the inlet (A11).

The average inlet microbial concentration for Campus B and Campus A differed by a factor of 10 (3 cells/ml vs 30 cells/ml), suggesting microbial differences in source water between the two campuses. Little variability was observed in live and total cells counts or the BQW values across Campus B (stations B15 and B16) and in the incoming water (stations B13 and B14, see Table S1). A comparison of the two testbeds (Figure S1) revealed that Campus A had lower levels of residual disinfectant (monochloramine) in the drinking water distribution systems than Campus B, presumably due to water ages of up to 100 h in the pumped storage system at the inlet of Campus A as well as roof storage for high-rise student accommodation near station A15. Lower levels of residual disinfectant on Testbed A may have promoted significant bacterial growth in the bulk water and biofilms as reported by FCM and enzymatic measurements of bacterial growth (BQW) compared to conditions on Testbed B (Table S1). Similar findings have been reported in the literature (Ling et al. 2018), where FCM measurements of water in premise plumbing systems showed a large increase in biomass before and after a week-long campus closure (10^3^ cells/ml and 7.8×10^5^ cells/ml, respectively) that also correlated with a decay of free residual chlorine (1.2 mg/L to ∼ 0.2 mg/L).

However, biofilm re-growth is not exclusively steered by changes in the levels of secondary disinfectants, and resistant biofilm-associated taxa belonging to *Pseudomonadaceae* and *Rhodospirillaceae* have been described as processing monochloramine during stagnation periods and utilizing it as a source of nutrients (Bédard et al. 2016). Monochloramine has been found to select for resistant bacterial populations in drinking water systems like *Legionella*, *Escherichia*, and *Geobacter* in a lab-scale system, and *Mycobacterium*, *Sphingmonas*, and *Coxiella* in a full-scale system (Chiao et al. 2014). Stanish et al (2016) also noticed a prevalence of microbes involved with nitrification and iron-cycling in chloramine-treated drinking water distribution systems (Stanish et al. 2016).

### 3.2 Composition of microbial communities in biofilms

Between February 2017 and August 2018, a total of 64 samples were collected, sequenced, and compared to characterize the microbial community structure of biofilms in the studied testbeds. Based on their origin (Testbed A or B), all biofilm samples were differentiated into four groups: MPB (MPB_A, n= 9; MPB_B, n = 2) and YSB (YSB_A, n = 38 resulted from 5 rounds of sampling; YSB_B, n = 15 resulted from 2 rounds of sampling). After rarefaction, 1,240 sequences were retained per sample at species level (97% sequence identity), which resulted in 708 and 491 OTUs in MPB from Testbed A and B, respectively; and 559 and 267 OTUs in YSB for A and B, respectively (Figure 1a). A total of 158 OTUs were found to be present in all the biofilm samples across the two Testbeds and were therefore recognized as the core community. The core community in both MPB and YSB comprised primarily of sequences identified as these four classes (with the range of relative abundance), Alphaproteobacteria (26 - 44%), Gammaproteobacteria (8 - 21%), Betaproteobacteria (3 - 28%), and Actinobacteria (2 - 17%) (Figure 1b; Table 2).

**Figure 1.**
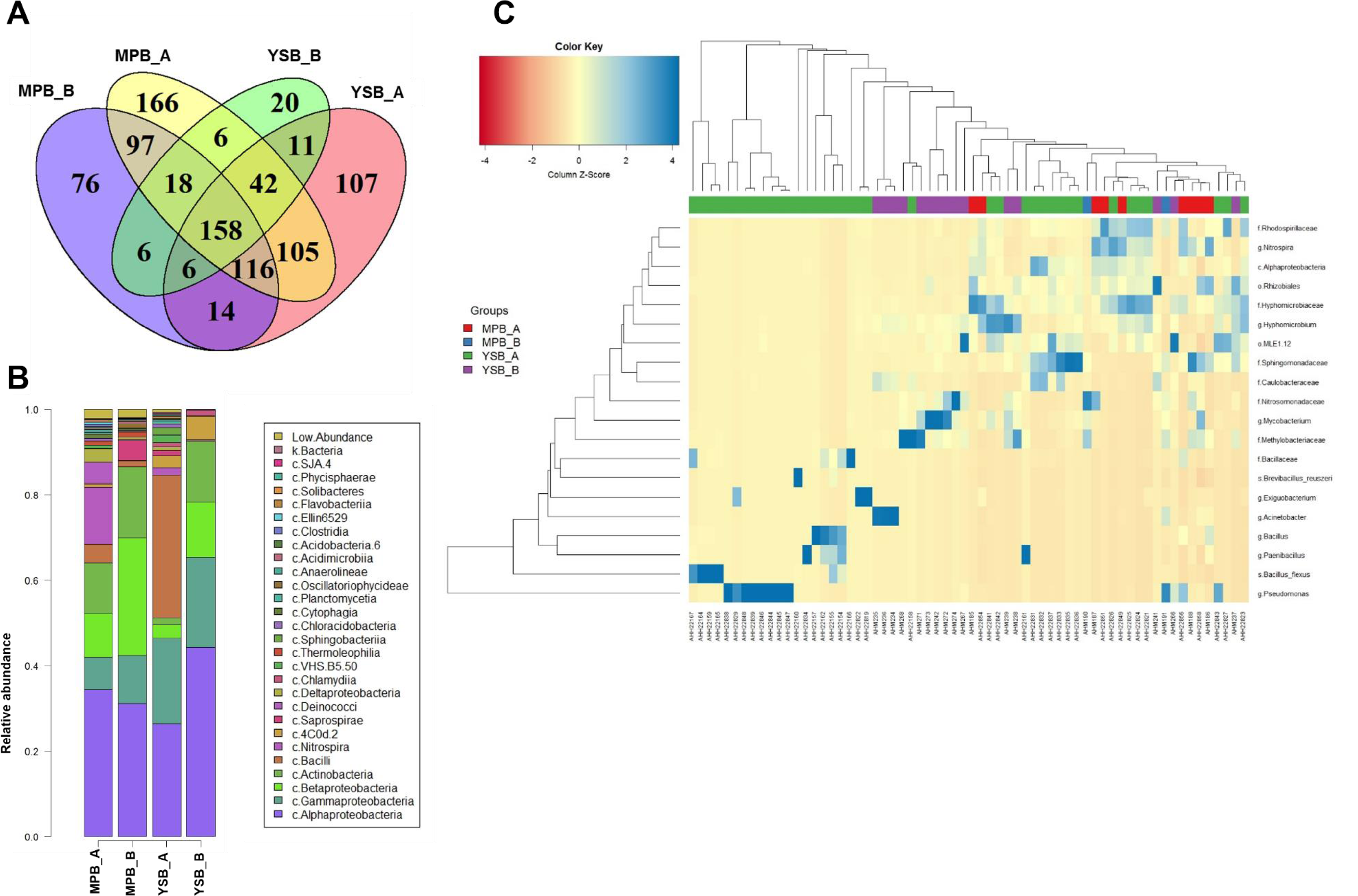
Overview of bacterial microbial community structure in the mature pipe biofilm (MPB) and young sensor biofilm (YSB) at two testbeds. a) Venn diagram of common OTUs in the four different biofilm categories, mature pipe biofilm from campus A (MPB_A, n=9) and B (MPB_B, n=2), young sensor biofilm from testbed A (YSB_A, n=38) and B (YSB_B, n=15); b) Relative abundance of bacterial community obtained from seven YSB at class level); c) heatmap showing the 20 most different taxa at OTU level in biofilm samples based on relative abundance.

**Table 2.** Average relative abundance of the ten most common core OTUs in mature pipe biofilms and young sensor biofilms from Testbeds A and B

| ID | Phylum | Class | Order | Family | Genus | MPB_A<br>(n=9) | MPB_B<br>(n=2) | YSB_A<br>(n=38) | YSB_B<br>(n=15) |
| --- | --- | --- | --- | --- | --- | --- | --- | --- | --- |
| 1 | Actinobacteria | Actinobacteria | Actinomycetales | Mycobacteriaceae | Mycobacterium | 0.51 | 2.46 | 0.38 | 12.23 |
| 2 | Proteobacteria | Alphaproteobacteria | Rhizobiales | Hyphomicrobiaceae | Hyphomicrobium | 2.82 | 0.04 | 2.09 | 6.80 |
| 3 | Proteobacteria | Alphaproteobacteria | Rhizobiales | Hyphomicrobiaceae |  | 8.42 | 0.30 | 2.87 | 1.44 |
| 4 | Proteobacteria | Alphaproteobacteria | Rhizobiales | Methylobacteriaceae |  | 0.58 | 6.39 | 1.49 | 11.79 |
| 5 | Proteobacteria | Alphaproteobacteria | Rhizobiales |  |  | 4.35 | 0.57 | 1.30 | 2.97 |
| 6 | Proteobacteria | Alphaproteobacteria | Rhodospirillales | Rhodospirillaceae |  | 4.98 | 1.62 | 2.09 | 0.21 |
| 7 | Proteobacteria | Betaproteobacteria | Burkholderiales | Comamonadaceae |  | 1.35 | 9.17 | 0.23 | 0.18 |
| 8 | Proteobacteria | Betaproteobacteria | Nitrosomonadales | Nitrosomonadaceae |  | 2.40 | 9.56 | 0.02 | 7.30 |
| 9 | Proteobacteria | Gammaproteobacteria | Pseudomonadales | Moraxellaceae | Acinetobacter | 0.19 | 1.79 | 0.02 | 12.26 |
| 10 | Proteobacteria | Gammaproteobacteria | Pseudomonadales | Pseudomonadaceae | Pseudomonas | 0.64 | 4.95 | 15.10 | 0.27 |

In all samples, YSB samples tended to show significantly lower diversity evidenced by smaller Hill numbers (both first order and second order) (Fig. S2) than their MPB counterparts (with mean ^2^D = 6.81, 15.56 for YSB and MPB, respectively) with a p-value = 0.001. This suggests that the bacterial communities of mature biofilms (> 14 years) were much more diverse than that of younger biofilms (< 2 years) for both Testbeds. Our finding agrees with previous studies in showing that microbial communities become more complex with biofilm age (Douterelo et al. 2019, Wang et al. 2014).

#### 3.2.1 The common families in all the biofilm samples

The most common families in all the biofilm samples were *Pseudomonadaceae*, *Methylobacteriaceae*, *Nitrosomonadaceae*, *Mycobacteriaceae*, Moraxellaceae, *Hyphomicrobiaceae*, *Commamonadaceae*, *Rhizobiales*, and *Rhodospirillaceae* (Table 2). Families with higher relative abundance in the MPB were also present in YSB from the same campus (Table 2). In contrast, the average relative abundance of *Pseudomonadaceae* decreased with biofilm age on Campus A (15.10% for YSB_A vs 0.64% for MPB_A) but increased on Campus B (0.27% vs 4.95% for YSB_B and MPB_B, respectively).

*Mycobacterium* showed a higher relative abundance of 12.23% for YSB_B and 2.46% for MPB_B as compared to 0.38% and 0.51% for YSB_A and MPB_A (Table 2). The observation of such a high abundance of Mycobacteria in the DWDS with high concentrations of monochloramine is consistent with previous studies conducted in the USA by Waak et al. (2019). It could indicate a survival mechanism for replication of *Mycobacterium* in water distribution networks with bactericidal conditions (Delafont et al. 2014, Loret and Dumoutier 2019, Taylor et al. 2000, Vaerewijck et al. 2005).

*Methylobacteriaceae* were found at higher relative abundance in biofilms from Testbed B (11.79% for YSB_B and 6.39% for MPB_B) than Testbed A (1.49% and 0.58%). This family comprises of a large group of bacteria within the order *Rhizobiales*, most of which are facultative anaerobes and able to grow with one-carbon compounds as sources of energy, fix nitrogen, produce pink carotenoid pigments, and stimulate plant growth (Szwetkowski and Falkinham III 2020). Their presence in Testbed B agrees with previous findings showing that a switch in secondary disinfectant from chlorine to monochloramine enriched the growth of *Methylobacteriaceae* (Zhang 2012). The high relative abundance of both *Mycobacteria* and *Methylobacteria* in biofilms of Testbed B with higher monochloramine was also consistent with results reported by Waak et al. (2019), in which a positive Spearman rank correlation has been found between the copy-number-normalized fractions of *Mycobacteria* and *Methylobacteria* in the biofilms of the chloraminated DWDS.

Similarly, the family *Nitrosomonadaceae* showed higher relative abundance under high monochloramine conditions on Campus B (7.30% for YSB and 9.56% for MPB) when compared to Campus A (0.02% for YSB_A and 2.40% for MPB_A). *Nitrosomonadaceae* (betaproteobacteria class) are ammonia oxidizers that play an important role in drinking water networks (Cruz et al. 2020, Kitajima et al. 2020, Pérez et al. 2019).

In contrast, Rhodospirillaceae were found to be more prevalent in biofilms on Testbed A (2.09% for YSB_A and 4.98% for MPB_A) compared to Testbed B (0.21% for YSB_B and 1.62% for MPB_B). This family is known for its common presence in aquatic environments, such as stagnant water in pipe systems (Baldani et al. 2014). It thus suggests some degree of water stagnation on Testbed A.

Figure 1c shows a heat map of the 20 taxa that differed the most among the four sets of biofilm samples in terms of relative abundance. There was a much lower abundance of Nitrospir (0.11% and 0.16% for MPB and YSB, respectively) and Bacilli (1.40% and 0.26%, respectively) on Campus B compared to Campus A (Nitrospira: 13.30% and 1.79%; and Bacilli: 4.42% and 33.46% for MPB and YSB, respectively). Younger biofilms showed a comparably higher relative bundance of Gammaproteobacteria (20.10% and 21.01% for Campus A and B, respectively) compared to their mature counterparts (7.57% and 11.25% for Campus A and B).

Genus *Pseudomonas* (within the Gammaproteobacteria class) was found at significantly higher relative abundance (an average relative abundance of 56.2% for YSB_A16_139d) in young biofilms at Station A16. Species including *Pseudomonas alcaligenes*, *Pseudomonas stutzeri*, and *Pseudomonas viridiflava* (but not the human pathogenic *P. aeruginosa)* were found in YSB after 36 -140 days. These data suggest their active role in colonizing the sensor surfaces that is consistent with prior pure culture studies (Ruinelli et al. 2017, Sommer et al. 2017, Suzuki et al. 2013).

The relative abundance table at order level (data not shown) further shows that Legionellales were found at low relative abundance in MPB and YSB from both Testbeds (average of 0.04%, 0.96%, and 0.46% for MPB_A, MPB_B, and YSB_A, respectively); they were completely absent from all 15 YSB_B samples. The complete absence of gene markers for *Legionella* spp., one of the most relevant opportunistic waterborne pathogens in engineered water systems, indicates the effectiveness of high monochloramine levels to suppress those bacteria within Testbed B. Similarly, Waak et al (2018) reported that *Legionella* spp were not detected by qPCR in the water-main biofilms of a chloraminated system in the USA, but were frequently detected in the in Norway where no residual chemical disinfectant is used, based on cut pipe sections during routine maintenance (Waak et al. 2018). Thus, maintenance of an appropriate level of chloramine residual throughout the campus is essential in suppressing the proliferation of *Legionella* in DWDS.

### 3.3 Factors shaping bacterial communities in MPB

NMDS analysis showed a clear temporal difference in the bacterial communities of MPB samples collected before October 31^st^, 2017 (thereafter referred as Phase I) and after October 2017 (Phase II) (p = 0.001; Figure 2a). The bioenv function (in the vegan package) identified biofilm age (p = 0.001) as the key environmental variable with the maximum (rank) correlation with community dissimilarities (Figure 2B), affecting the occurrence of both *Acinetobacter lwoffii* (a potential opportunistic pathogen) and *Bacillus selenatarsenatis*. *Bacillus selenatarsenatis* has been known for its ability to utilize arsenate, selenite, or nitrate as electron acceptors under aerobic conditions or ferment carbohydrates like glucose, lactate and pyruvate under anaerobic conditions (Yamamura et al. 2007).

**Figure 2.**
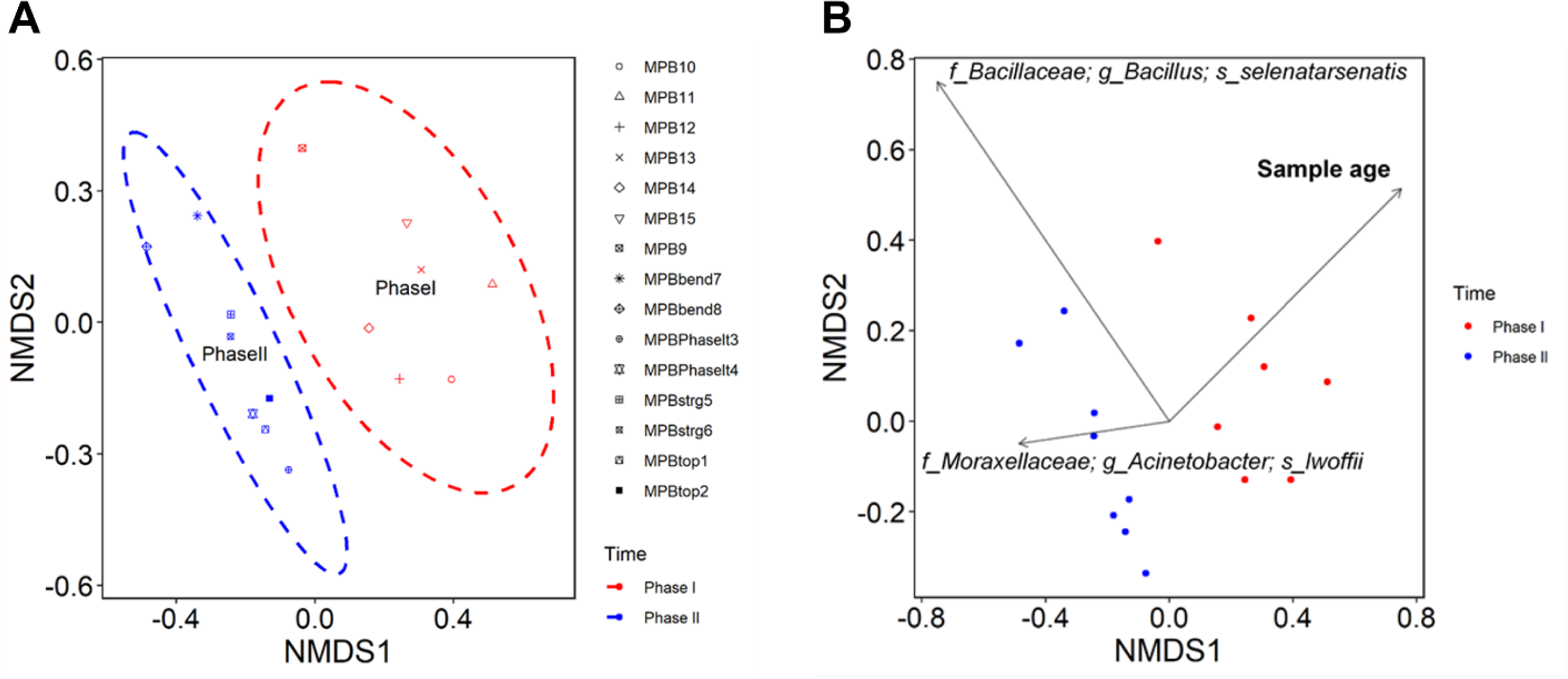
Nonmetric multidimensional scaling (NMDS) of OTU level compositions of MPB samples and the significant environmental variables with highest Pearson correlation with community dissimilarities plotted using the envfit function of VEGAN. a) Temporal change of bacterial composition within MPB samples; b) Vectors and significant taxa with an average abundance greater than 1%. Note: Phase I, mature pipe biofilm (MPB) samples collected prior to November 1, 2017; phase II, MPB samples collected after November 1, 2017. The dash lines indicate 95% confidence level.

MPB samples collected in Phase I showed a relatively higher biodiversity than those of Phase II (Figure. S3). Conductivity and sample age have been shown to have a positive correlation with the inverse Simpson index (^2^D, second order Hill number) with a correlation coefficient value of 0.48, and 0.45, respectively (p < 0.1) according to the regression of ^2^D against the collected limited environmental variables (Figure. S3). This could also be explained from the long-term monitoring of conductivity data. A notable decrease was noticed in the average conductivity on Campus A from 187 ± 28 µS/cm (July to October 2017, Phase I) to 162 ± 19 µS/cm (November 2017 to Jun 2018, Phase II) and moderate decrease in conductivity on Campus B from 296 ± 28 µS/cm in Phase I to 284 ± 36 µS/cm in Phase II during the same time window. This change of MPB over time is consistent with earlier findings described by Ling et al. (2016) who observed that temporal variation has a more significant contribution to the biofilm community difference than other factors, including infrastructural distribution system characteristics (i.e. pipe material), pipe age, or residual disinfectant type.

### 3.4 Correlations between water physiochemical factors and MPB bacterial communities

Canonical correlation analysis (CCA), which uses analysis of variance tests based on Bray-Curtis distance matrices, revealed conductivity, sample age, and pipe diameter as important environmental variables influencing the bacterial community structure for MPB, while water pressure was less significant (Figure 3a). This agrees with the NMDS analysis showing a temporal effect on nascent community structure in biofilms (see Figure 2).

**Figure 3.**
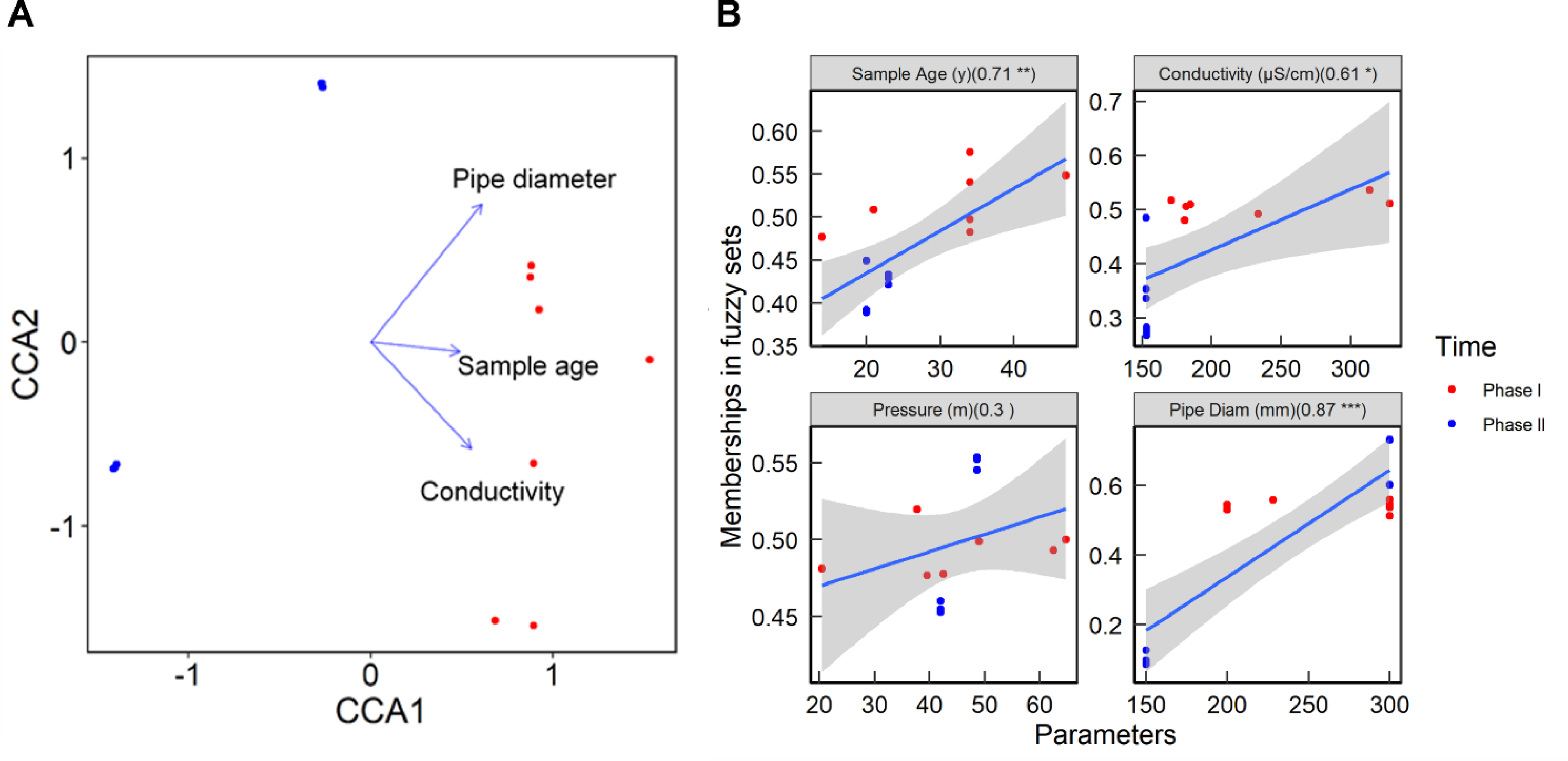
Environmental factors influencing the community structure of MPB samples identified by a) canonical correspondence Analysis (CCA) and b) fuzzy set ordination (FSO). The number on the top panel of each plot indicates the correlation coefficient with permutated probability (· p<0.1, * p<0.05, **p<0.01, ***<0.001). The similarity index is calculated based on Horn method and the dissimilarity index is calculated by (1-similarity index). Note: Phase I, mature pipe biofilm (MPB) samples collected prior to November 1, 2017; phase II, MPB samples collected after November 1, 2017.

Similarly, the FSO analysis (Fig. 3b) suggested that conductivity, biofilm age (Table 2), and pipe diameter correlated with changes in the bacterial community of MPB (correlation coefficients 0.61, 0.71, and 0.87 with p-values of 0.018, 0.003, and 0.001, respectively. There was little influence of water pressure. The observed relationship between conductivity and biofilm community agrees with an earlier study that reported progressive long-term salinity change promoted the assembly of new communities (Logares et al. 2013).

### 3.5 Dynamics of YSB development

A total of 7 YSB samples were collected (5 from Campus A with biofilm ages of 36 - 202 days and 2 from Campus B with biofilm ages between 326 and 405 days, Figure S1d) during sensor maintenance operations. Bacilli (within the phylum of Firmicutes) initially colonized the biofilm in the first 2 months, followed by Proteobacteria and Cyanobacteria, for both stations (A14 and A16) investigated on Campus A (Figure 4). The community profiles from the same station were further analyzed and compared using the multidimensional scaling (MDS) algorithm. The sequence data separated into two distinct data sets for YSB communities clustered by different biofilm ages measured at Stations A14 (65 vs 138 days; Figure 5a) and A16 (35 vs 139 days; Figure 5b). The plots include outcomes for each different sensor material separately. The permutational ANOVA p-value is 0.001 in each case; thus, it is readily apparent that biofilm age had a much greater influence on the biofilm community than the sensor surface material.

**Figure 4.**
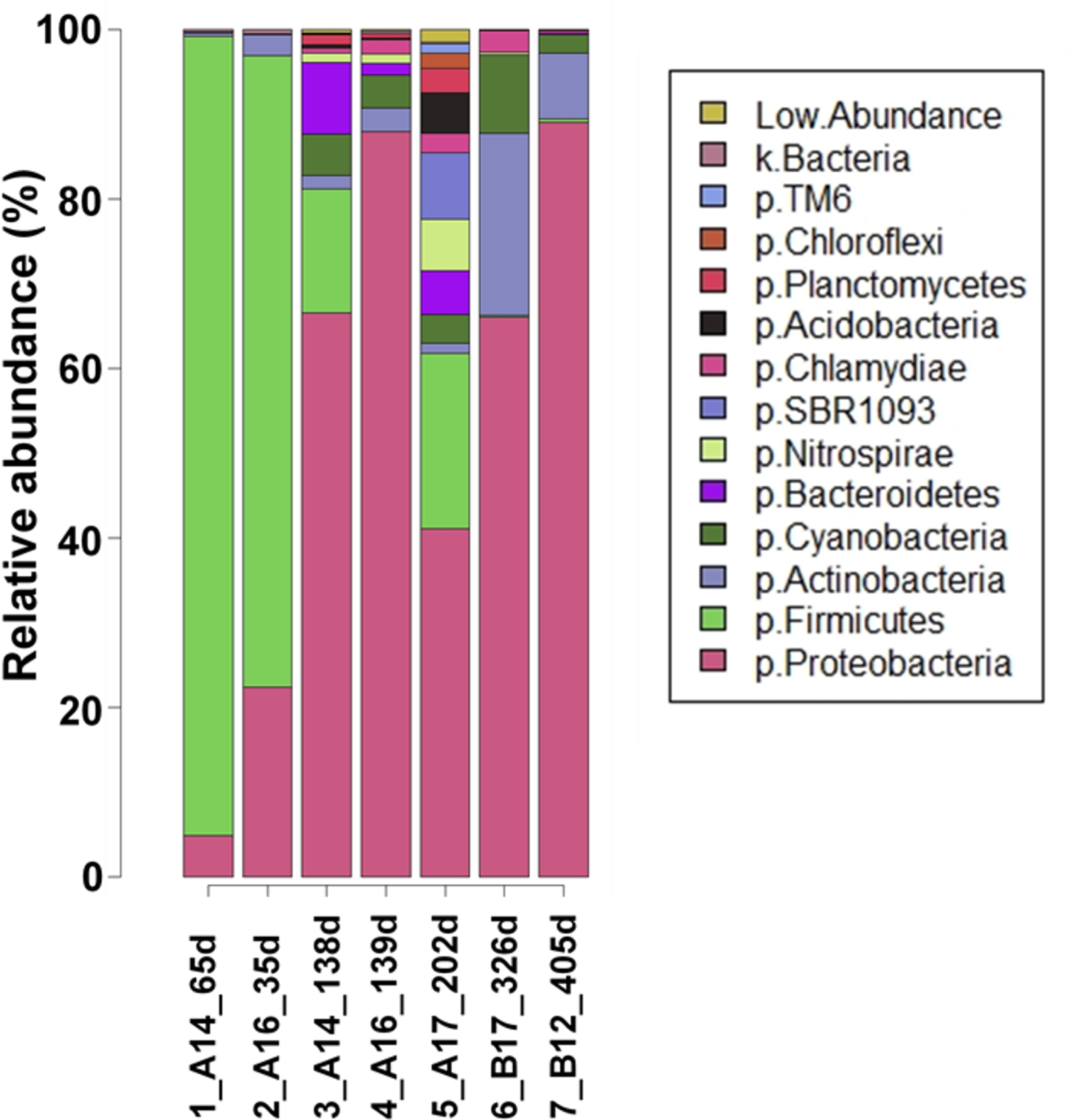
Overview of bacterial microbial community structure at phylum level in seven young sensor biofilm (YSB) samples

**Figure 5.**
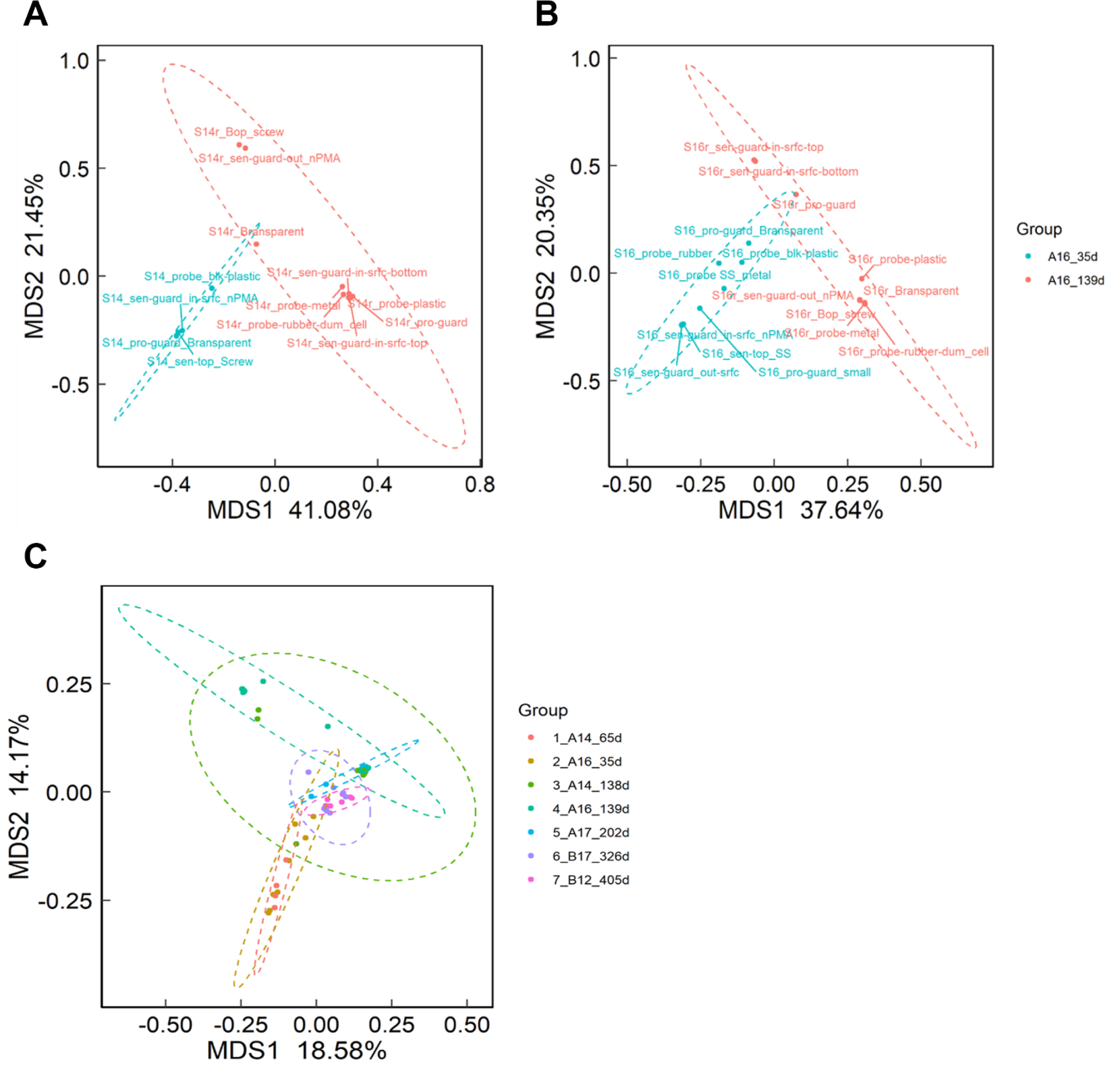
Clustering of young sensor biofilms (YSBs) by biofilm age at OTU level: a) Biofilm sampled two times from A14 (Permanova, p = 0.001); b) Biofilm sampled two times at A16 (Permanova, p = 0.001); c), Clustering of all the YSB samples by sampling location and age. The dash lines indicate 95% confidence level. Permutational multivariate analysis of variance (PERMANOVA) was applied to assess the differences in bacterial community structure among samples and it shows that significant community difference is seen from YSB of different biofilm age from the same station A14, but also A16. The sample name shown in the MPD plot indicates different part of the sensor material.

Figure 5c shows MDS analyses for all seven YSB samples and further supports the observed clustering of microbial community based on sample location, thus indicating that both location and biofilm age were stronger drivers for bacterial community composition than sensor surface type. Station A17 (biofilm age of 202 days) had the highest diversity based on the calculated Hill numbers (^2^D=16.16, FigS4a). These analyses confirm that diversity (and therefore microbiome complexity) increased with age for the young biofilms on Campus A. A similar change was also noticed for the YSB on Campus B.

### 3.6 Correlations between water physiochemical factors and YSB bacterial communities

CCA plots were generated to explore the relationship of environmental variables with the community difference. Among these variables, Figure 6 shows that pH, water age, conductivity, DTM, and YSB sample age all contributed to the community differences among YSB samples.

**Figure 6.**
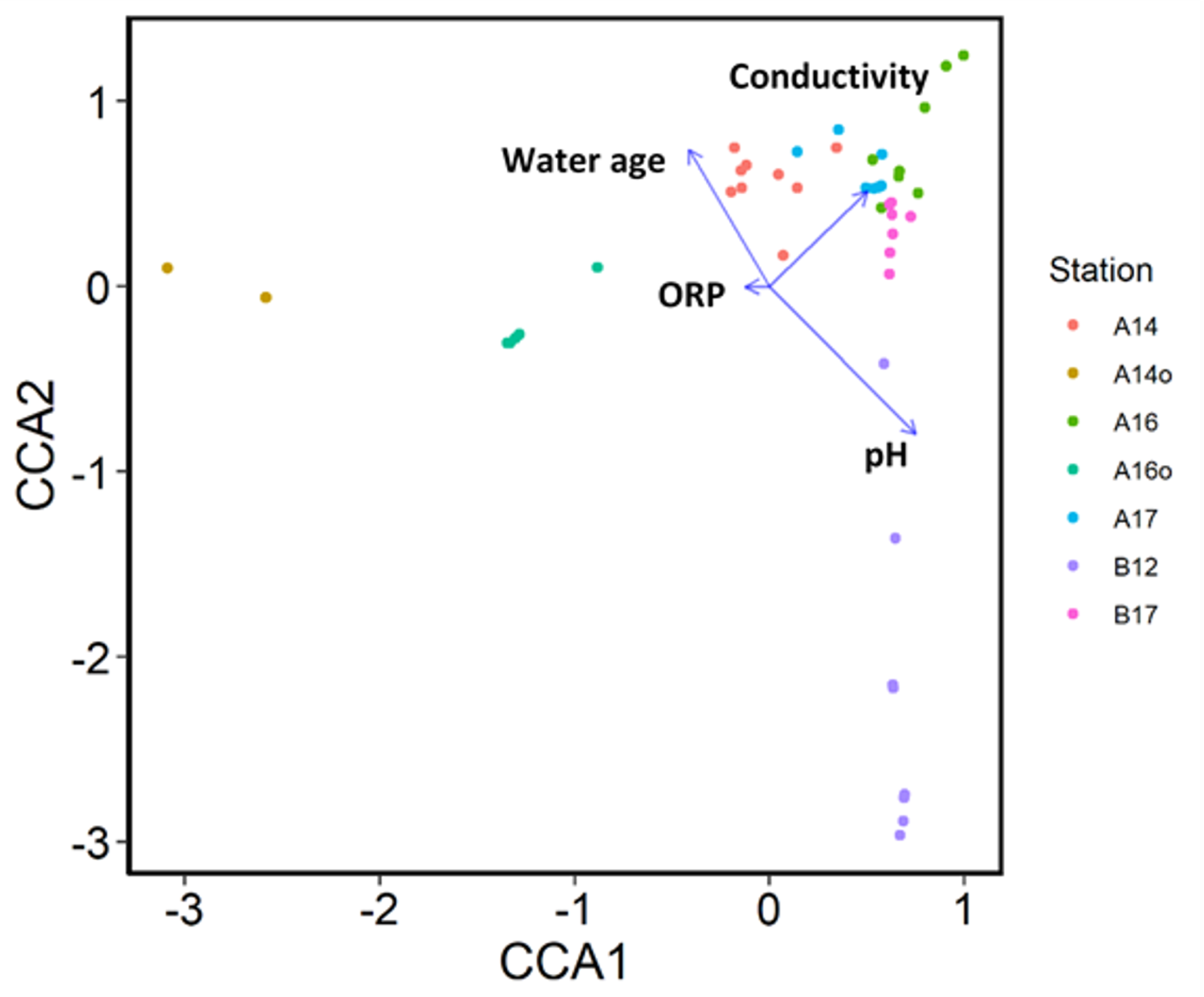
Correlation of water age, conductivity, ORP and pH with bacterial community among all YSB samples based on canonical correlation analysis (CCA). CCA which uses analysis of variance tests based on Bray-Curtis distance matrix revealed pH, conductivity, water age as well as ORP as important environmental variables influencing the bacterial community structure for YSB. Note: A14o and A16o represent the first sampling date (June 15th, 2017) of YSB at stations A14 and A16 on campus A since installation, and A14 and A16 indicate the second sampling date (October 31st and November 1st 2017, respectively).

Multivariate analysis based on FSO showed that pH and sample age had a significant influence on the YSB community difference with a correlation coefficient greater than 0.7 (Figure S5). The higher the membership value in fuzzy sets, the greater the impact on the YSB community difference. The results are visualized by correlating the dissimilarity between fuzzy set membership against original values (at reported levels of significance). The influence of sample age and pH was significantly higher than that of sulphate, chloride, monochloramine, ammonia, and nitrate concentrations (0.6 < r < 0.7) in the bulk water. This was followed by water age, TN, TIC, pressure, TOC and pipe diameter (0.5 < r < 0.6) on both campuses. DTM, nitrite, TC, conductivity, orthophosphorus and ORP had a relatively weak impact (r < 0.5) on the community composition in the testbeds, based on these FSO analyses.

### 3.7 The impact of biofilm on water quality and potential biofilm indicator microorganisms

The relationship between YSB and various water quality parameters was investigated at all stations in the two Testbeds. Most of the parameters included agreed well with the reported city water quality parameters, except for a spike in PO_4_-P values at two stations on two different days, which coincided with a clear baseline shift in DTM readings (Figure S6). Stations A16 and B17 registered a significantly higher average concentration of PO_4_-P (0.38 mg/L) than the other stations. This value was also higher than the reported concentration of city water (0.017 mg/L). YSB was sampled at these monitoring stations on the days the shift in baseline PO_4_-P occurred and again after 91 (for A16) and 76 days (for B17), when there was a significant decrease in PO_4_-P level in the bulk water phase (Table S4).

An increase in the DTM has been previously proposed as an indirect approach to monitoring the biofilm development (Flemming et al. 2011). Placing the water quality sensor probes in the stagnant bulk water during preliminary experiments also showed a rapid DTM shift from the turbidity probe of the water quality sensor (data not shown). Here, we found a gradual increase in the baseline for only two of the turbidity sensors, Station A16 and B17 (Figure S6a and S6b), while offline turbidity analyses showed a 0 FNU reading from all the stations tested. This suggests that the biofilm development rates were faster compared to those in the remainder of the two testbeds. In other words, these turbidity sensors with an increasing baseline indicate possible water stagnation in regions around Stations A16 and B17 (these have been confirmed using hydraulic models of the respective campus networks). Further work is needed to understand the co-occurrence of DTM and PO_4_-P in operational DWDS.

In summary, this study proposes that the microbial community dynamics observed in YSB could be used as a proxy to assess the wider microbiome of MPB in inaccessible places and the DWDS at large. A portion of the bacterial community overlapped in both young and mature biofilms at various locations. Some of the identified taxa may act as sentinels for biofilm growth. Non-tuberculous *Mycobacteria* have been known for their survival strategy in water networks or under harsh conditions and their ability to grow slowly by taking advantage of an amoebae host for persistence and replication (Delafont et al. 2014). In addition, the *Acinetobacter* spp. (member of the family Moraxellaceae) inhabited the YSB on Campus B at an average relative abundance of 12.3%, despite consistently high concentrations of monochloramine (Table 2).

*Acinetobacter* spp. were also identified in YSB samples from both campuses using the plate count method with R2A medium, thus proving their viability. The genus *Acinetobacter*, containing several species known to cause significant burdens of disease as nosocomial pathogens, was recently described as a leading factor in the horizontal transfer of antimicrobial resistance genes in biofilm environments (Almasaudi 2018, Joshi and Litake 2013, Khan et al. 2017) and cells are known to develop resistance to disinfection and desiccation (Wong et al. 2017).

## 4. Conclusions

Our findings provide novel insights into the factors shaping biofilm community dynamics within two operational campus drinking water distribution systems that supply a total of 80,000 individuals in an urban setting:

- Higher levels of residual monochloramine were found on Campus B. These conditions appear to suppress regrowth of biomass in the bulk water (and sensor biofilms) compared to conditions on Campus A. However, this situation may also promote resistant *Acinetobacter* spp. and Mycobacteria in both mature and young biofilms on Campus B. *Pseudomonas* spp. together with *Bacillus* spp. are likely the earlier colonizers leading to the initial formation of biofilm (YSB) on Campus A with reduced monochloramine conditions and relatively higher water age.
- Differences in the community structure were evident when mature biofilm samples were grouped according to their collection time, suggesting the strong impact of temporal effects and water chemistry. However, further studies should be conducted on the young biofilms aged between 1 and 5 years to allow for insights into the biofilm development associated with their interaction with localized bulk water.
- Conductivity, sample age, and pipe diameter correlated to the MPB community based on both CCA and FSO analysis. MPB showed higher biodiversity than YSB and the ^2^D of YSB positively correlates with conductivity, but negatively with ORP or nitrite. The microbial community dynamics observed in YSB could be used as a proxy to assess the microbiome of relatively inaccessible (underground) MPB.
- The age of bulk water as well as biofilm together with their respective location in the DWDS had significantly higher influence on the community composition than the substrate for YSB.Biofilm age (p = 0.001) was identified as the factor showing the maximum correlation with community dissimilarities by FSO analysis. In addition, the influence of pH and sample age on the YSB community is greater than the amounts of sulphate, chloride, monochloramine, ammonia, and nitrate in the bulk water, followed by water age, TN, TIC, pressure, TOC and pipe diameter. DTM, nitrite, TC, conductivity, orthophosphorus and ORP seem to have a relative weak impact on the community composition in the testbeds under investigation.
- Significant levels of orthophosphate were detected in YSB samples at two stations (A16 and B17) and associated with higher levels of stagnation based on long-term DTM. Further work is required to verify their interaction and subsequent importance in the DWDS.

## Acknowledgements

This research was supported by the National Research Foundation (NRF), Singapore through the Singapore-MIT Alliance for Research and Technology (SMART) Center of Environmental Sensing and Modeling. This research was also supported by the NRF, Prime Minister’s Office, Singapore under its Campus for Research Excellence and Technological Enterprise (CREATE) program, and by the Singapore Ministry of Education through a RCE award to Singapore Centre for Environmental Life Sciences Engineering (SCELSE).

We thank the staff at the facilities management office of the two campuses for providing access to the networks and information about operational changes within the studied periods. We are also grateful to Ami Preis (Xylem) during the initiation, operation, and support of the online monitoring platform for these two testbeds. We thank de Sessions Paola Florez for help with the sequencing service, and Eric Alm and Gu Xiaoqiong for their discussion of the Venn diagram. The authors would like to thank Eric Dubois Hill for his invaluable help with the visualization of the data.

## Contribution

CD and AJW conceived and planned the study. CD and CM collected and processed the samples and conducted the physico-chemical analysis. CD generated the figures, conducted the bioinformatical analysis and wrote the initial draft of the manuscript. SW, US, JRT and ML provided valuable feedback and helped shape the discussion, analysis and overall narrative. All authors contributed by proofreading and editing the manuscript.

## Data availability

All data supporting the findings of this study are available from the corresponding author upon request.

## References

Alberdi, A. and Gilbert, M.T.P. (2019) A guide to the application of Hill numbers to DNA-based diversity analyses. Mol. Ecol. Resour. 19, 804–817.

Allaire, M., Wu, H. and Lall, U. (2018) National trends in drinking water quality violations. Proc. Natl. Acad. Sci. U S A 115, 2078–2083.

Allen, M., Preis, A., Iqbal, M. and Whittle, A.J. (2013) Water distribution system monitoring and decision support using a wireless sensor network, Hawaii, USA.

Almasaudi, S.B. (2018) *Acinetobacter* spp. as nosocomial pathogens: epidemiology and resistance features. Saudi J. Biological Sci. 28, 586–596.

Anderson, M.J. (2014-2017) Permutational multivariate analysis of variance (PERMANOVA), p. 07841, John Wiley & Sons, Ltd, New Zealand.

Ashbolt, N.J. (2015) Microbial contamination of drinking water and human health from community water systems. Curr. Environ. Health Rep. 2, 95–106.

Baldani, J.I., Videira, S.S., Teixeira, K.R.d.S., Reis, V.M., Oliveira, A.L.M.d., Schwab, S., Souza, E.M.d., Pedraza, R.O., Baldani, V.L.D. and Hartmann, A. (2014) The Prokaryotes. E., R., E.F., D., Lory S., S.E. and F., T. (eds), Springer, Berlin, Heidelberg.

Bédard, E., Prévost, M. and Déziel, E. (2016) *Pseudomonas aeruginosa* in premise plumbing of large buildings. MicrobiologyOpen 5, 937–956.

Boyce, R.L. and Ellison, P.C. (2001) Choosing the best similarity index when performing fuzzy set ordination on binary data. J. Vegetation Sci. 12, 711–720.

Buse, H.Y., Lu, J., Lu, X., Mou, X. and Ashbolt, N.J. (2014) Microbial diversities (16S and 18S rRNA gene pyrosequencing) and environmental pathogens within drinking water biofilms grown on the common premise plumbing materials unplasticized polyvinylchloride and copper. FEMS Microbial. Ecol. 88, 280–295.

Carminati, M., Turolla, A., Mezzera, L., Mauro, M.D., Tizzoni, M., Pani, C., Zanetto, F., Foschi, J. and Antonelli, M. (2020) A self-powered wireless water quality sensing network enabling smart monitoring of biological and chemical stability in supply systems. Sensors 20, s20041125.

Chao, A., Gotelli, N.J., Hsieh, T.C., Sander, E.L., Ma, K.H., Colwell, R.K. and Ellison, A.M. (2014) Rarefaction and extrapolation with Hill numbers: a framework for sampling and estimation in species diversity studies. Ecol. Monogr. 84, 45–67.

Chiao, T.-H., Clancy, T.M., Pinto, A., Xi, C. and Raskin, L. (2014) Differential resistance of drinking water bacterial populations to monochloramine disinfection. Environ. Sci. Technol. 48, 4038–4047.

Cruz, M.C., Woo, Y., Flemming, H.-C. and Wuertz, S. (2020) Nitrifying niche differentiation in biofilms from full-scale chloraminated drinking water distribution system. Water Res. 176, 115738.

Delafont, V., Mougari, F., Cambau, E., Joyeux, M., Bouchon, D., Héchard, Y. and Moulin, L. (2014) First evidence of amoebae-mycobacteria association in drinking water network. Environ. Sci. Technol. 48, 11872–11882.

Rossman, L. A. (2000). EPANET 2 USERS MANUAL. U. S. E. P. A. Water supply and Water Resources Division. Cincinnati, OH.

Douterelo, I., Boxall, J.B., Deines, P., Sekar, R., Fish, K.E. and Biggs, C.A. (2014) Methodological approaches for studying the microbial ecology of drinking water distribution systems. Water Research 65, 134–156.

Douterelo, I., Dutilh, B.E., Arkhipova, K., Calero, C. and Husband, S. (2020) Microbial diversity, ecological networks and functional traits associated to materials used in drinking water distribution systems. Water Res. 173, 115586.

Douterelo, I., Sharpe, R.L., Husband, S., Fish, K.E. and Boxall, J.B. (2019) Understanding microbial ecology to improve management of drinking water distribution systems. WIREs Water 6, e01325.

Fang, W., Hu, J.Y. and Ong, S.L. (2009) Influence of phosphorus on biofilm formation in model drinking water distribution systems. J. Appl. Microbiol. 106, 1328–1335.

Flemming, H.-C., Tamachkiarowa, A., Klahre, J. and Schmitt, J. (1998) Monitoring of fouling and biofouling in technical systems. Water Sci. Technol. 38, 291–298.

Flemming, H.-C., Wingender, J. and Szewzyk, U. (2011) Biofilm Highlights, Springer, Heidelberg, Dordrecht, London, New York.

Flemming, H.-C., Wingender, J., Szewzyk, U., Steinberg, P., Rice, S.A. and Kjelleberg, S. (2016) Biofilms: an emergent form of bacterial life. Nat Rev Micro 14, 563–575.

Flemming, H.-C. and Wuertz, S. (2019) Bacteria and archaea on Earth and their abundance in biofilms. Nature Rev. Microbiol. 17, 247–260.

Henne, K., Kahlisch, L., Brettar, I. and Höfle, M.G. (2012) Analysis of structure and composition of bacterial core communities in mature drinking water biofilms and bulk water of a citywide network in Germany. Appl. Environ. Microbiol. 78, 3530–3538.

Hull, N.M., Ling, F., Pinto, A.J., Albertsen, M., Jang, H.G., Hong, P.-Y., Konstantinidis, K.T., Lechevallier, M., Colwell, R.R. and Liu, W.-T. (2019) Drinking Water Microbiome Project: Is it Time? Trends in Microbiology 27, 670.

Janknecht, P. and Melo, L.F. (2003) Online biofilm monitoring. Reviews in Environ. Sci. Bio/Technol. 2, 269–283.

Joshi, S.G. and Litake, G.M. (2013) Acinetobacter baumannii: an emerging pathogenic threat to public health. World J. Clin. Infect. Dis. 3, 25–36.

Kasahara, S., Maeda, K. and Ishikawa, M. (2004) Influence of phosphorus on biofilm accumulation in drinking water distribution systems. Water Sci. Technol. 4, 389–398.

Khan, H.A., Baig, F.K. and Mehboob, R. (2017) Nosocomial infections: epidemiology, prevention, control and surveillance. Asian Pacific J. Tropical Biomedicine 7, 478–482.

Kitajima, M., Cruz, M.C., Williams, R.B., Wuertz, S. and Whittle, A.J. (2020) Microbial abundance and community composition in biofilms on in-pipe sensors in drinking water distribution systems. Sci. Total Environ. In review.

Ling, F., Hwang, C., LeChevallier, M.W., Andersen, G.L. and Liu, W.-T. (2016) Core-satellite populations and seasonability of water meter biofilms in a metropolitan drinking water distribution system The ISME J 10, 582–595.

Ling, F., Whitaker, R., LeChevallier, M.W. and Liu, W.-T. (2018) Drinking water microbiome assembly induced by water stagnation. The ISME J 12, 1520–1531.

Liu, G., Bakker, G.L., Li, S., Vreeburg, J.H.G., Verberk, J.Q.J.C., Medema, G.J., Liu, W.T. and Dijk, J.C.V. (2014) Pyrosequencing reveals bacterial communities in unchlorinated drinking water distribution system: an integral study of bulk water, suspended solids, loose deposits, and pipe wall biofilm. Environ. Sci. Technol. 48, 5467–5476.

Logares, R., Lindström, E.S., Langenheder, S., Logue, J.B., Paterson, H., Laybourn-Parry, J., Rengefors, K., Tranvik, L. and Bertilsson, S. (2013) Biogeography of bacterial communities exposed to progressive long-term environmental change. ISME J 7, 937–948.

Loret, J.-F. and Dumoutier, N. (2019) Non-tuberculous mycobacteria in drinking water systems: A review of prevalence data and control means. Int. J. Hygiene & Environ. Health 222.

Miller, C.S., Baker, B.J., Thomas, B.C., Singer, S.W. and Banfield, J.F. (2011) EMIRGE: reconstruction of full-length ribosomal genes from microbial community short read sequencing data. Genome Biology 12, R44.

Miller, C.S., Handley, K.M., Wrighton, K.C., Frischkorn, K.R., Thomas, B.C. and Banfield, J.F. (2013) Short-read assembly of full-length 16S amplicons reveals bacterial diversity in subsurface sediments. PLoS ONE 8, e56018.

Ong, S.H., Kukkillaya, V.U., Wilm, A., Lay, C., Ho, E.X.P., Low, L., Hibberd, M.L. and Nagarajan, N. (2013) Species identification and profiling of complex microbial communities using shotgun illumina sequencing of 16S RNA amplicon sequences. PLoS ONE 8, e60811.

Pérez, M.F.L., Cárdenas1, J.A., Leon1, A.J.M. and Susa, M.S.R. (2019) Changes on eubacteria 338i, gamma-, betaproteobacteria in biofilms from drinking water systems according to operational conditions Environ. Processes 6, 85–106.

PUB (2016) Managing the water distribution network with a Smart Water Grid. Smart Water 1, 4.

Raskin, L. and Nielsen, P.H. (2019) Editorial overview: Integrating biotechnology and microbial ecology in urban water infrastructure through a microbiome continuum viewpoint. Curr. Opin. Biotechnol. 57.

Reeslev, M., Nielsen, J.C. and Rogers, L. (2011) Assessment of the bacterial contamination and remediation efficacy after flooding using fluorometric detection. J. ASTM Internl. 8.

Roberts, D.W. (2009) Comparison of multidimensional fuzzy set ordination with CCA and DB-RDA. Ecology 90, 2622–2634.

Ruinelli, M., Blom, J. and Pothier, J.F. (2017) Complete genome sequence of *Pseudomonas viridiflava* cfbp 1590, isolated from diseased cherry in france. Genome Announc. 5, e00662–00617.

Schwake, D.O., Garner, E., Strom, O.R., Pruden, A. and Edwards, M.A. (2016) *Legionella* DNA makers in tap water coincident with a spike in Legionnaires’ disease in Flint, MI. Environ. Sci. Technol. Letters 3, 311–315.

Sommer, M., Xie, H. and Michel, H. (2017) Pseudomonas stutzeri as an alternative host for membrane proteins. Microb. Cell Factories 16, 157.

Stanish, L.F., Hull, N.M., Robertson, C.E., Harris, J.K., Stevens, M.J., Spear, J.R. and Pace, N.R. (2016) Factors influencing bacterial diversity and community composition in municipal drinking waters in the Ohio river basin, USA. PLoS ONE 11, e0157966.

Strathmann, M., Mittenzwey, K.-H., Sinn, G., Papadakis, W. and Flemming, H.-C. (2013) Simultaneous monitoring of biofilm growth, microbial activity and inorganic deposits on surfaces with an optical deposit sensor in-situ, on-line, in real time and non-destructively. Biofouling 29, 573–583.

Suzuki, M., Suzuki, S., Matsui, M., Hiraki, Y., Kawano, F. and Shibayama, K. (2013) Genome sequence of a strain of the human pathogenic bacterium *Pseudomonas alcaligenes* that caused bloodstream infection. Genome Announc. 1, e00919–00913.

Szwetkowski, K.J. and Falkinham III, J.O. (2020) *Methylobacterium* spp. as emerging opportunistic premise plumbing pathogens. Pathogens 9, 9020149.

Taylor, R.H., Falkinham, J.O., Norton, C.D. and LeChevallier, M.W. (2000) Chlorine, chloramine, chlorine dioxide, and ozone susceptibility of *Mycobacterium avium*. Appl. Environ. Microbiol. 66, 1702–1705.

Torondel, B., Ensink, J.H.J., Gundogdu, O., Ijaz, U.Z., Parkhill, J., Abdelahi, F., Nguyen, V.A., Sudgen, S., Gibson, W., Walker, A.W. and Quince, C. (2016) Assessment of the influence of intrinsic environmental and geographical factors on the bacterial ecology of pit latrines. Microb. Biotechnol. 9, 209–223.

Vaerewijck, M.J.M., Huys, G., Palomino, J.C., Swings, J. and Portaels, F. (2005) Mycobacteria in drinking water distribution systems: ecology and significance for human health FEMS Microbiol. Rev. 29, 911–934.

Waak, M.B., LaPara, T.M., Hallé, C. and Hozalski, R.M. (2018) Occurrence of *Legionella* spp. in water-main biofilms from two drinking water distribution systems. Environ. Sci. Technol. 52, 7630–7639.

Waak, M.B., LaPara, T.M., Hallé, C. and Hozalski, R.M. (2019) Nontuberculous Mycobacteria in two drinking water distribution systems and the role of residual disinfection. Environ. Sci. Technol. 53, 8563–8573.

Wang, H., Masters, S., Edwards, M.A., Falkinham, J.O.I. and Pruden, A. (2014) Effect of disinfectant, water age, and pipe materials on bacterial and eukaryotic community structure in drinking water biofilm. Environ. Sci. Technol. 48, 1426–1435.

Wong, D., Nielsen, T.B., Bonomo, R.A., Pantapalangkoor, P., Luna, B. and Spellberg, B. (2017) Clinical and pathophysiological overview of acinetobacter infections: a century of challenges. Clin. Microbiol. Rev. 30, 409–447.

Yamamura, S., Yamashita, M., Fujimoto, N., Kuroda, M., Kashiwa, M., Sei, K., Fujita, M. and Ike, M. (2007) *Bacillus selenatarsenatis* sp. nov., a selenate- and arsenate-reducing bacterium isolated from the effluent drain of a glass-manufacturing plant. Int. J. System. Evol. Microbiol. 57, 1060–1064.

Zhang, Y. (2012) Microbial Community Dynamics and Assembly: Drinking Water Treatment and Distribution, University of Tennessee, Knoxville.

