## Supplementary Tables and Figures for "Factors Shaping Young and Mature Bacterial Biofilm Communities in Two Drinking Water Distribution Networks"

A

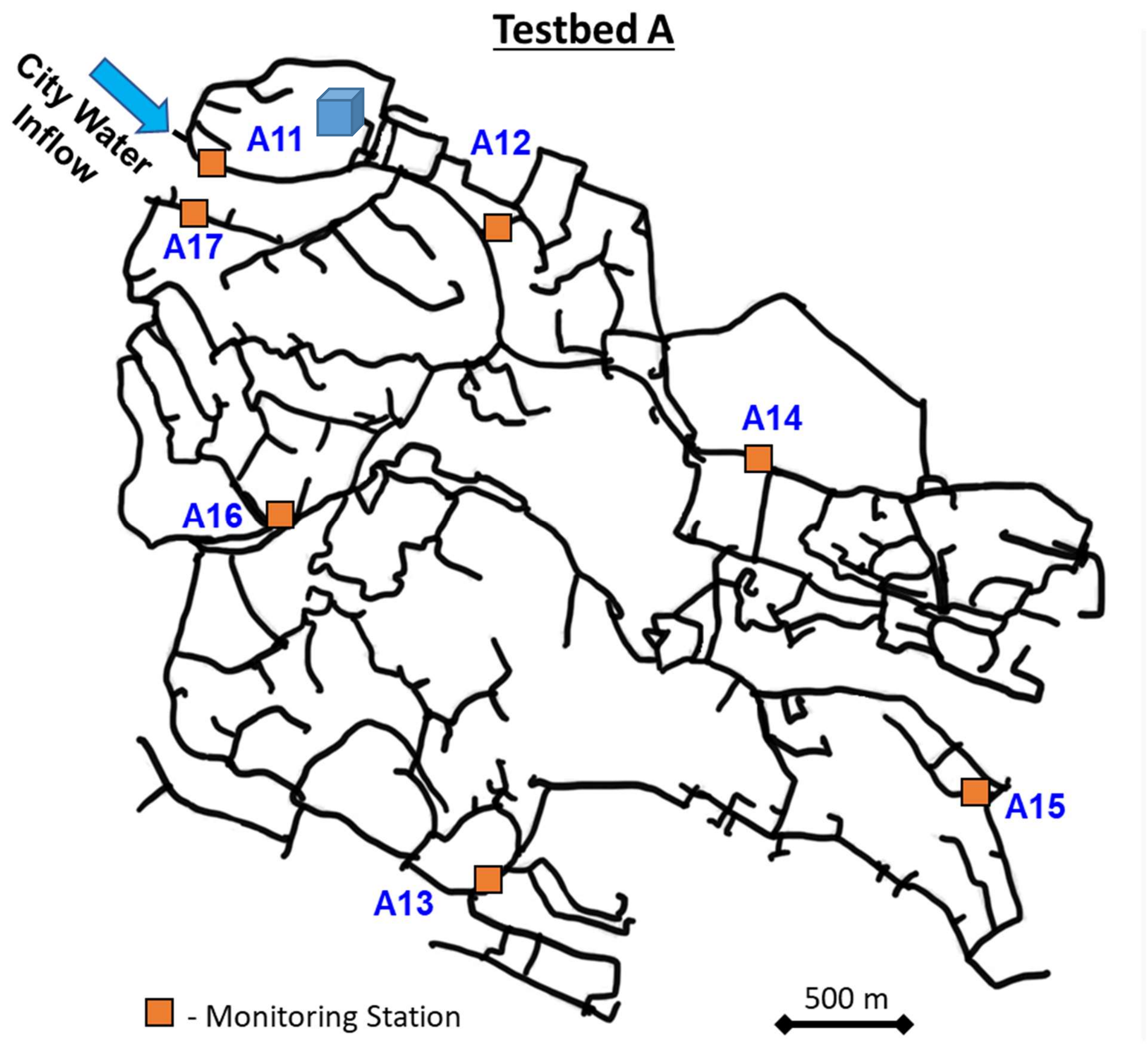

Testbed A
City Water Inflow
A11
A12
A17
A14
A16
A15
A13
- Monitoring Station
500 m

B

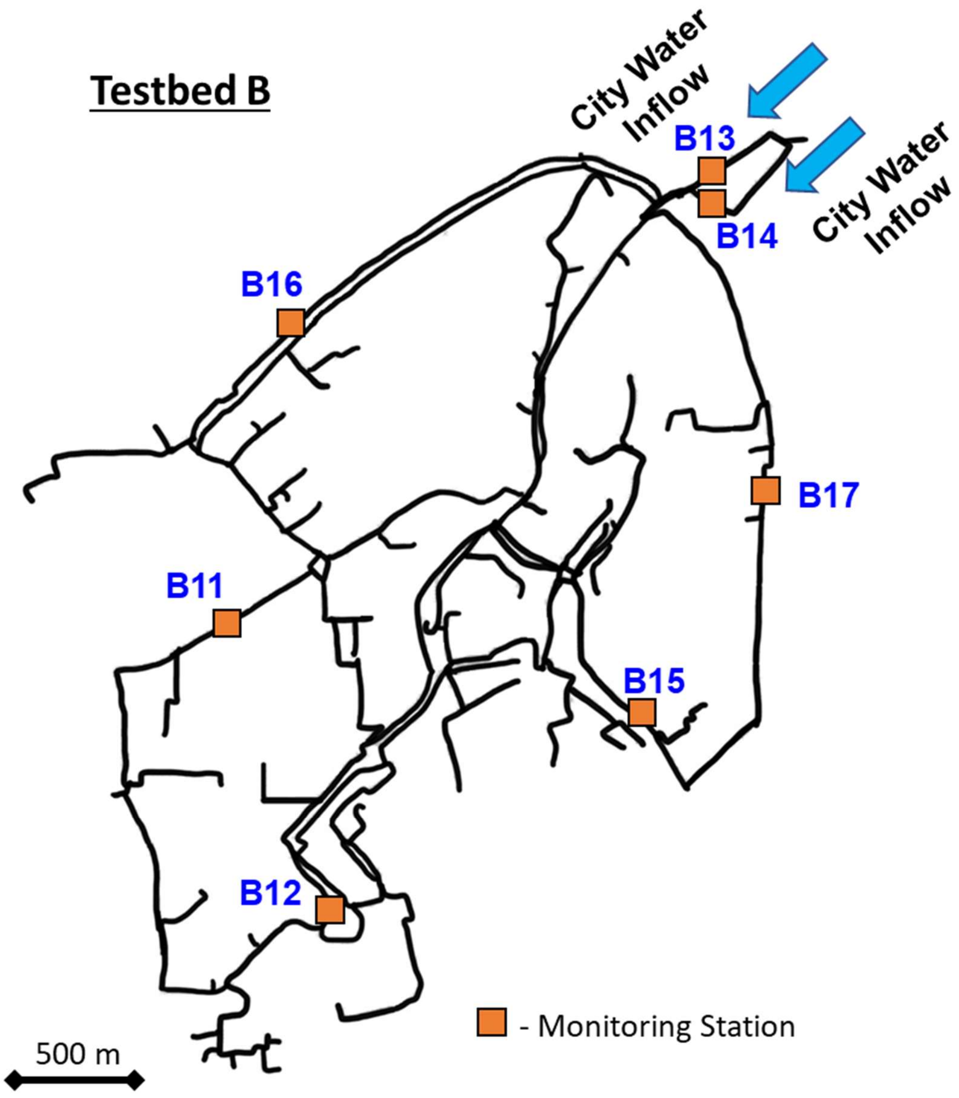

Testbed B
City Water Inflow
City Water Inflow
B13
B14
B16
B17
B11
B15
B12
- Monitoring Station
500 m

C

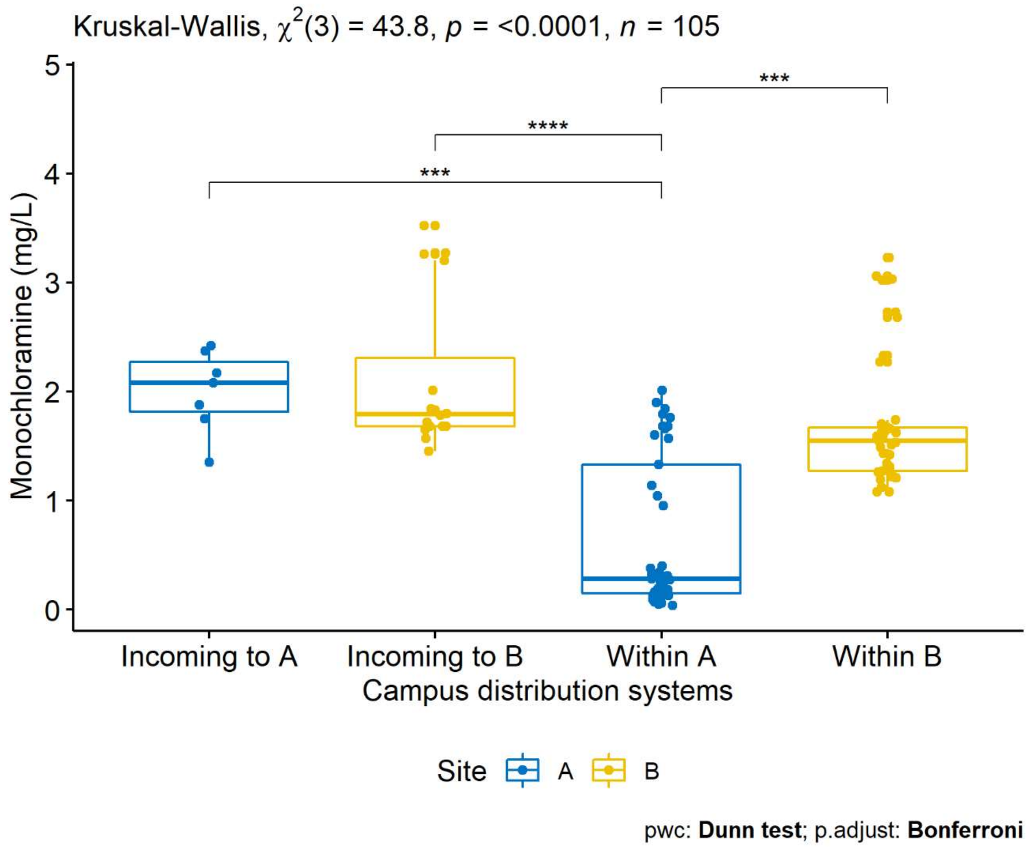

Kruskal-Wallis, $\chi^2(3)$ = 43.8, $p$ = <0.0001, $n$ = 105
Monochloramine (mg/L)
5
4
3
2
1
0
***
****
***
Incoming to A
Incoming to B
Within A
Within B
Campus distribution systems
Site
A
B
pwc: **Dunn test**; p.adjust: **Bonferroni**

D

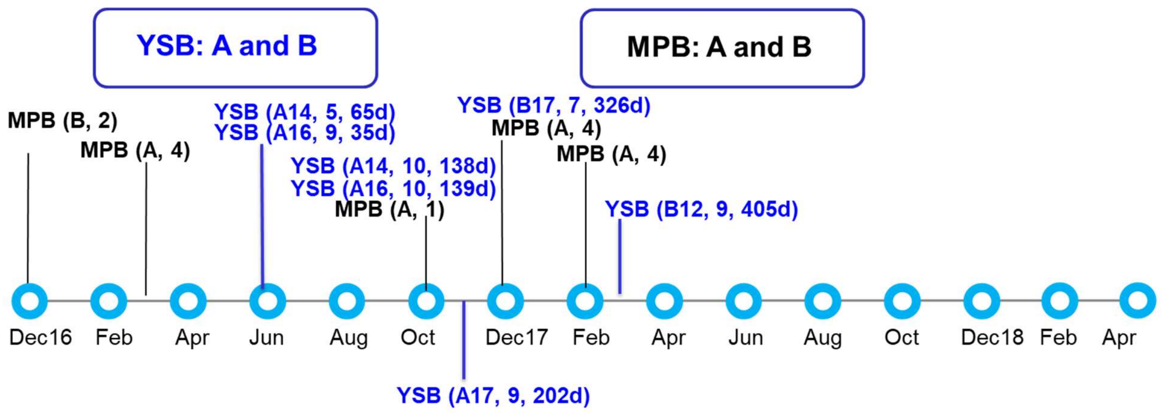


Figure S1. Campus testbeds for collection of biofilm samples. A) DWDS and sensor stations for Campus A (inlet at A11) serving about 40,000 individuals daily; B) DWDS and sensor stations for Campus B (inlet at B13 and B14) serving about 40,000 individual daily; C) Boxplot of the monochloramine concentration from two campus testbeds A and B during Feb 2017 to Dec 2019; D) Sampling timeline for young sensor biofilm (YSB) and mature pipe biofilm (MPB) from campus testbeds A and B. The time interval between two circles was 2 months. Sample ID explanation: MPB (B, 2) refers to two samples collected and sequenced from campus B; YSB (A14, 5, 65 d) means that five samples with a biofilm age of 65 d were collected from station A14. The sampling of MPB has been divided into Phase I (samples taken before October 2017) and Phase II (samples taken after October 2017).

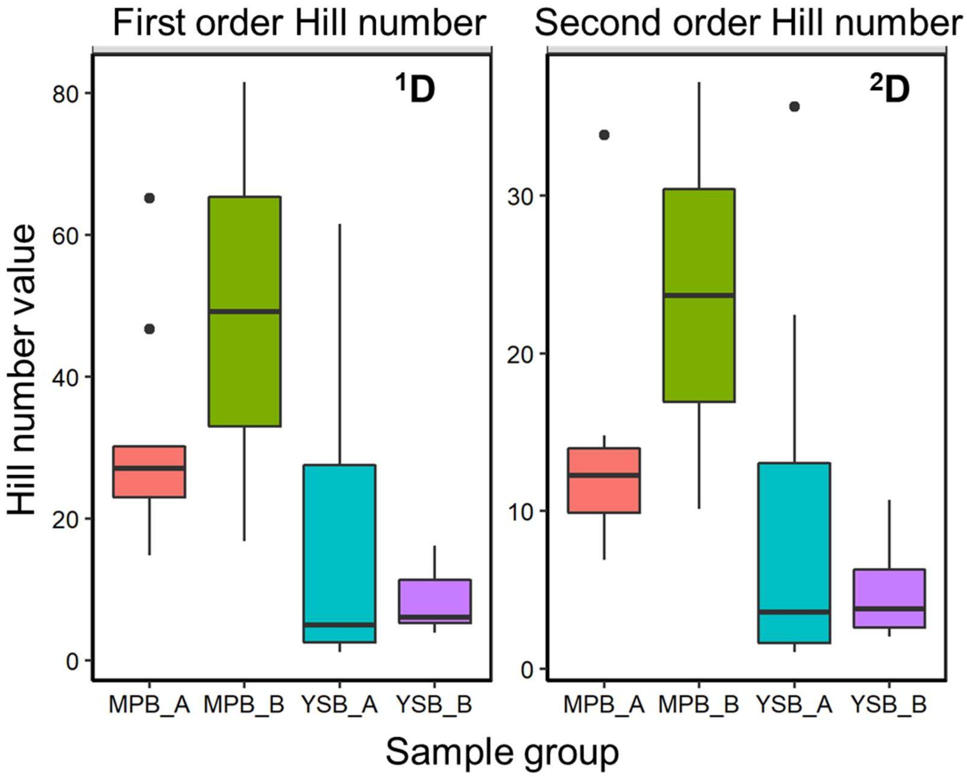


Figure S2. Boxplot of Hill numbers from mature pipe biofilms (MPB) and young sensor biofilms (YSB) collected from two campuses A and B.

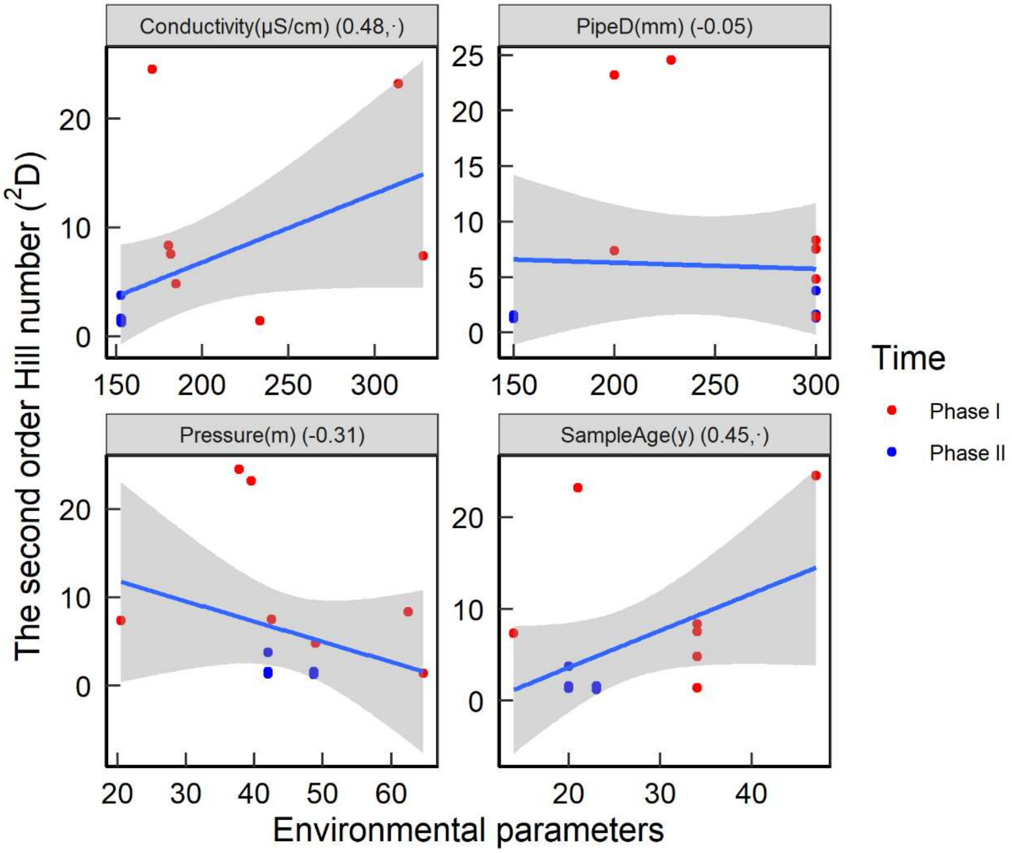


Figure S3. The second order Hill number of MPB samples as a function of conductivity, pipe diameter, pressure and pipe age (sample age). The number on the top panel of each plot indicates the correlation coefficient with permutated probability (· $p<0.1$, * $p<0.05$, ** $p<0.01$, *** $p<0.001$). None of the relationships are significant.

A

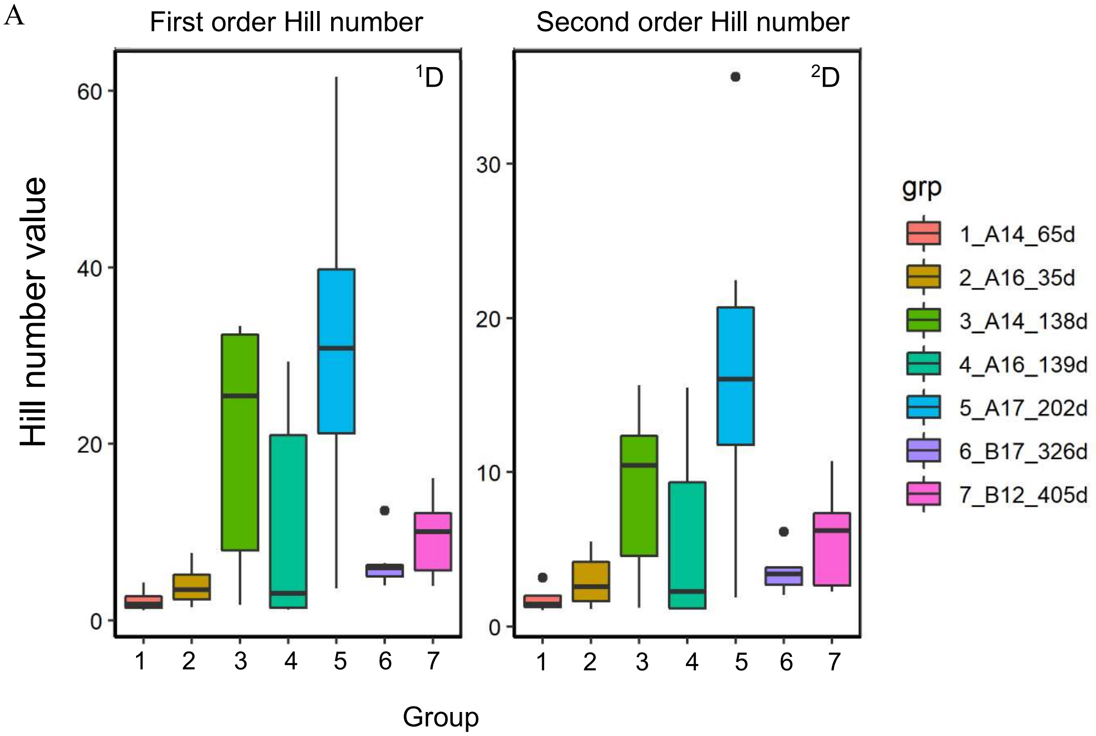
First order Hill number
Second order Hill number
$^{1}D$
$^{2}D$
Hill number value
Group
grp
1_A14_65d
2_A16_35d
3_A14_138d
4_A16_139d
5_A17_202d
6_B17_326d
7_B12_405d

B

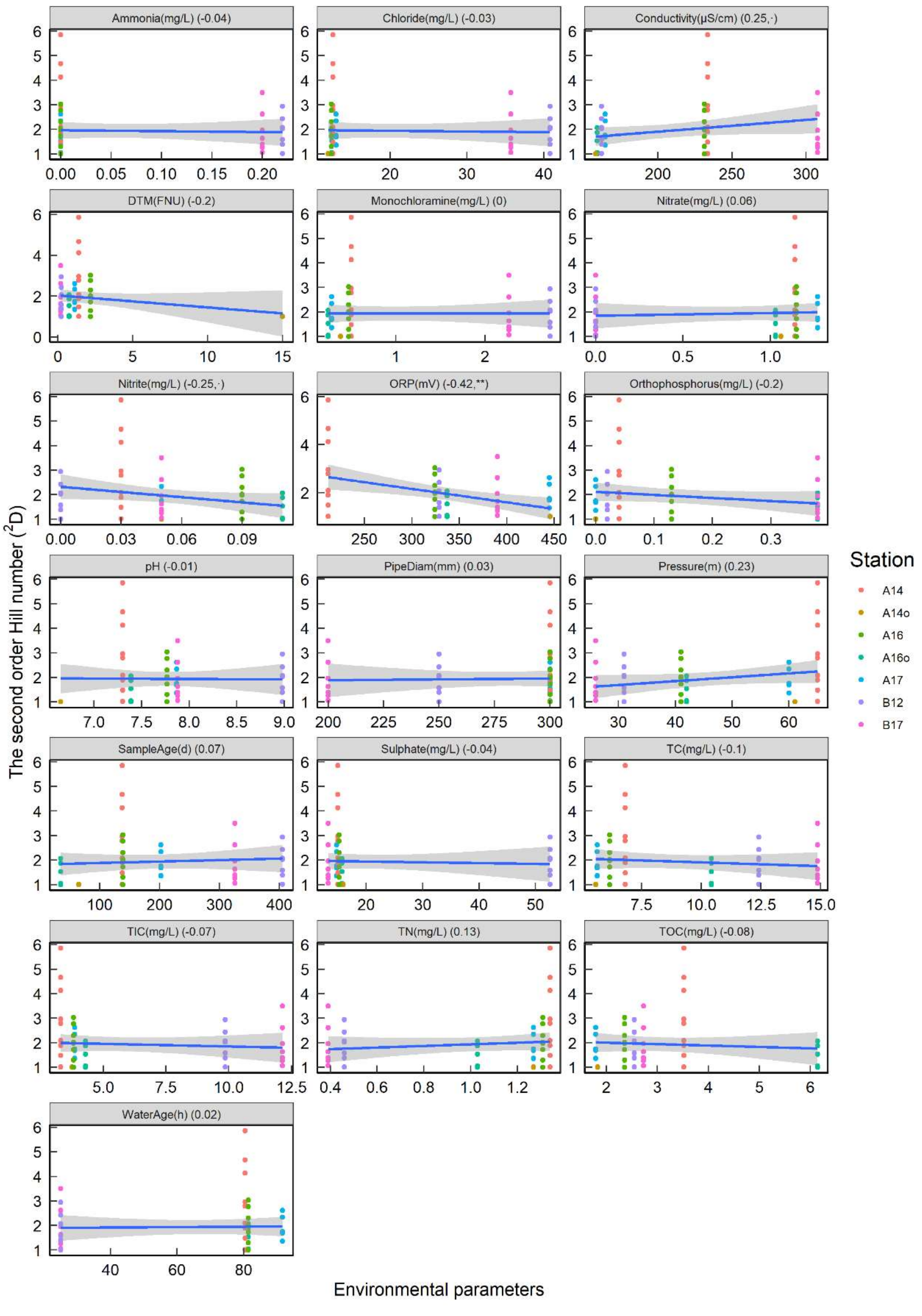

Ammonia(mg/L) (-0.04)
Chloride(mg/L) (-0.03)
Conductivity(µS/cm) (0.25,·)
DTM(FNU) (-0.2)
Monochloramine(mg/L) (0)
Nitrate(mg/L) (0.06)
Nitrite(mg/L) (-0.25,·)
ORP(mV) (-0.42,**)
Orthophosphorus(mg/L) (-0.2)
pH (-0.01)
PipeDiam(mm) (0.03)
Pressure(m) (0.23)
SampleAge(d) (0.07)
Sulphate(mg/L) (-0.04)
TC(mg/L) (-0.1)
TIC(mg/L) (-0.07)
TN(mg/L) (0.13)
TOC(mg/L) (-0.08)
WaterAge(h) (0.02)
The second order Hill number ($^2$D)
Environmental parameters
Station
A14
A14o
A16
A16o
A17
B12
B17

Fig S4. Hill numbers of young sensor biofilms (A) and the linear correlation (B) between individual environmental parameters and second order Hill number ($^2$D). Each scatter plot shows a different environmental variable, including ammonia, chloride, conductivity, DTM, monochloramine, nitrate, nitrite, ORP, orthophosphorus, pH, pipe diameter, pressure, sample age, sulphate, total carbon (TC), total inorganic carbon (TIC), total nitrogen (TN), total organic carbon (TOC), and water age. The number on the top panel of each plot indicates the correlation coefficient with permutated probability (· $p<0.1$, * $p<0.05$, **$p<0.01$, ***$<0.001$). Pearson's correlations are given, and significance is indicated (· $p<0.1$, * $p<0.05$, **$p<0.01$, ***$<0.001$) in the panel headings. Only ORP had a significant correlation with $^2$H. Samples A14o and A16o refer to the first round of YSB samples with biofilm ages of 65 d and 35 d, respectively. The shaded area indicates the confidence intervals around the mean.

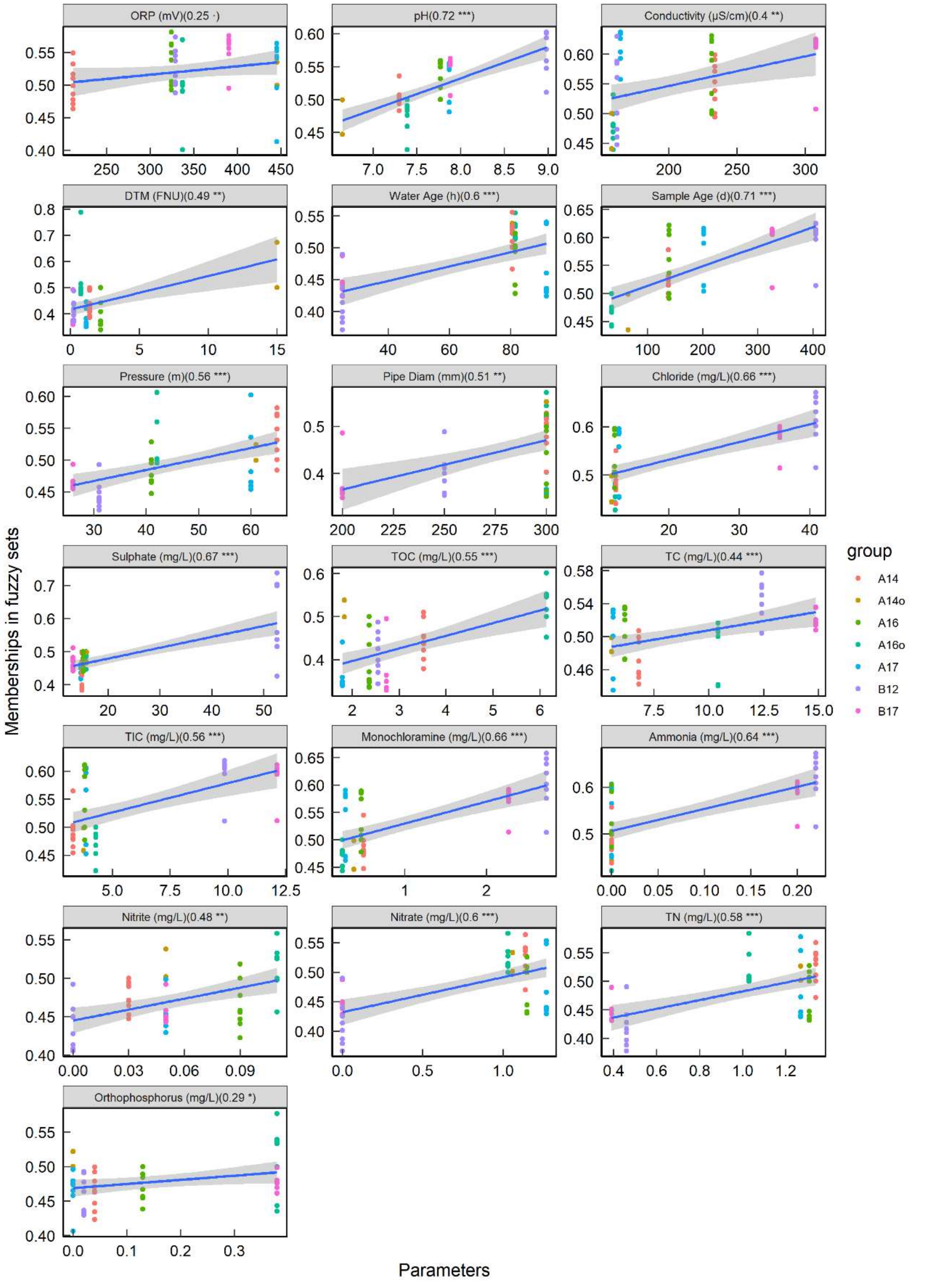
ORP (mV)(0.25 ·)
pH(0.72 ***)
Conductivity (μS/cm)(0.4 **)
DTM (FNU)(0.49 **)
Water Age (h)(0.6 ***)
Sample Age (d)(0.71 ***)
Pressure (m)(0.56 ***)
Pipe Diam (mm)(0.51 **)
Chloride (mg/L)(0.66 ***)
Sulphate (mg/L)(0.67 ***)
TOC (mg/L)(0.55 ***)
TC (mg/L)(0.44 ***)
TIC (mg/L)(0.56 ***)
Monochloramine (mg/L)(0.66 ***)
Ammonia (mg/L)(0.64 ***)
Nitrite (mg/L)(0.48 **)
Nitrate (mg/L)(0.6 ***)
TN (mg/L)(0.58 ***)
Orthophosphorus (mg/L)(0.29 *)
Memberships in fuzzy sets
Parameters
group
A14
A14o
A16
A16o
A17
B12
B17

Figure S5. Fuzzy set ordination (FSO) plot reflecting the membership function of environmental variables on the community variability (dissimilarity matrix) of YSB samples. The dissimilarity matrix of the biofilm community structure is calculated based on the distance index using Horn method (Roberts 2009, Torondel et al. 2016). For each of the specified variables, a fuzzy set ordination is calculated and the correlation between the original variable and the fuzzy set is reported. The significance of a particular variable is assessed by comparing a specified threshold p-value and the probability of obtaining a correlation between the data and fuzzy set. The results are visualised by producing a plot of fuzzy set against original values which is annotated with a correlation between them and a significance label. The number on the top panel of each plot indicates environmental variable (unit) and the correlation coefficient with permutated probability (· $p<0.1$, * $p<0.05$, **$p<0.01$, ***$<0.001$).

A

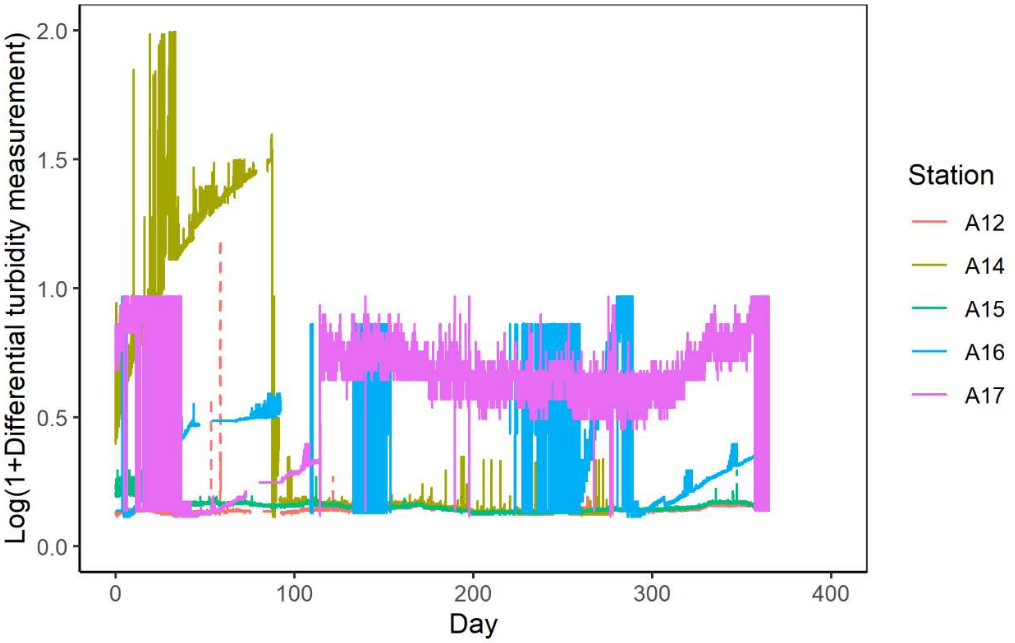


B

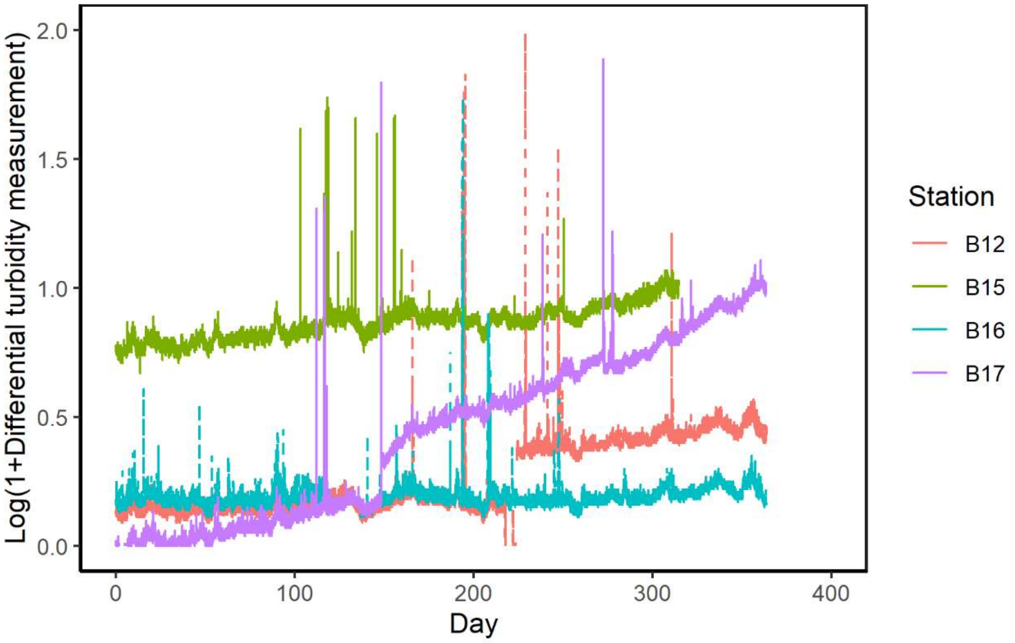


Figure S6. Biofilm accumulation as change in differential turbidity measurement (DTM) at testbeds from 1-Aug-17 (day 0) to 31-Jul-18 (day 364). A, testbed A; B, testbed B. The value of DTM is represented by the difference between the in-pipe turbidity sensor and the offline turbidity. The unit is FNU for DTM.

Supporting information

Table S1. Characterization of bulk water at selected stations from testbeds A and B

| Stn | Water Age (h) | Date | Monochloramine (mg/L) | Bacterial cell count (cells/ml) | | | | Live /total (%) | Particle count (n/ml) | Bactiquant water value* (BQW) |
|---|---|---|---|---|---|---|---|---|---|---|
| | | | | Live | Dead | Intermediate | Total | | | |
| A11 | 0 | 10-Apr-18 | 2.17 | 30 ± 23 | 1714 ± 393 | 23 ± 13 | 1767 ± 424 | 1.7 | 11938 ± 1815 | 9 |
| A12 | 85.4 | 9-Apr-18 | 0.40 | 58 ± 28 | 1050 ± 110 | 15 ± 9 | 1123 ± 130 | 5.2 | 11485 ± 1520 | 1509 |
| A15 | 164.1 | 11-Apr-18 | 0.36 | 65 ± 35 | 758 ± 272 | 25 ± 7 | 1040 ± 315 | 6.3 | 10160 ± 6021 | 1223 |
| A16 | 81.1 | 12-Apr-18 | 0.29 | 60 ± 5 | 745 ± 83 | 8 ± 6 | 813 ± 84 | 7.4 | 11198 ± 701 | 3330 |
| B13 | 0 | 16-Apr-18 | 1.87 | 3 ± 3 | 200 ± 13 | 3 ± 3 | 207 ± 8 | 1.6 | 2473 ± 140 | 27 |
| B14 | 0 | 17-Apr-18 | 1.05 | 0 ± 0 | 284 ± 72 | 0 ± 0 | 284 ± 72 | 0.0 | 7332 ± 1290 | 26 |
| B15 | 20-30 | 18-Apr-18 | 1.25 | 5 ± 0 | 528 ± 110 | 5 ± 5 | 538 ± 117 | 0.9 | 6820 ± 926 | 37 |
| B16 | 20-30 | 19-Apr-18 | 0.41 | 0 ± 0 | 275 ± 25 | 2 ± 2 | 277 ± 28 | 0.0 | 12888 ± 1638 | 21 |

Notes: Inlet stations: A11 for campus A; B13 and B14 for campus B. Particle count refers to organic or inorganic particulate matter larger than 1 µm. *Results are shown as dimensionless BactiQuant Water value (BQW) and reflect the number of bacteria based on enzyme activity.

**Table S2. Pipe and bulk water characteristics for mature pipe biofilm (MPB) samples from Testbeds A and B**

| ID | Sample name (Phase_Stn_material_location) | Pipe age (years) | Conductivity (µS/cm) | Pressure (m) | Pipe diameter (mm) | Sampling date | Campus |
|---|---|---|---|---|---|---|---|
| 1 | II_16N_DICL_top* | 20 | 152.89 | 41.99 | 300 | 13-Dec-17 | A |
| 2 | II_16N_DICL_top | 20 | 152.89 | 41.99 | 300 | 13-Dec-17 | A |
| 3 | II_16N_DICL_bottom | 20 | 152.89 | 41.99 | 300 | 13-Dec-17 | A |
| 4 | II_16N_DICL_bottom | 20 | 152.89 | 41.99 | 300 | 13-Dec-17 | A |
| 5 | II_21_DI_straight | 23 | 153.05 | 48.66 | 150 | 23-Feb-18 | A |
| 6 | II_21_DI_straight | 23 | 153.05 | 48.66 | 150 | 23-Feb-18 | A |
| 7 | II_21_DI_bend | 23 | 153.05 | 48.66 | 150 | 23-Feb-18 | A |
| 8 | II_21_DI_bend | 23 | 153.05 | 48.66 | 150 | 23-Feb-18 | A |
| 9 | I_14_DICL_coupon | 34 | 233.58 | 64.71 | 300 | 31-Oct-17 | A |
| 10 | I_13_DICL_coupon | 34 | 180.44 | 62.46 | 300 | 23-Mar-17 | A |

| | | | | | | | |
|---|---|---|---|---|---|---|---|
| 11 | I_15_DI_coupon | 47 | 171.03 | 37.73 | 228 | 25-Mar-17 | A |
| 12 | I_16_DICL_coupon | 34 | 181.70 | 42.47 | 300 | 21-Mar-17 | A |
| 13 | I_17_DICL_coupon | 34 | 184.76 | 48.98 | 300 | 17-Mar-17 | A |
| 14 | I_15_DIPL_coupon^ | 14 | 328.26 | 20.50 | 200 | 14-Dec-16 | B |
| 15 | I_16_DICL_coupon_ | 21 | 313.88 | 39.50 | 200 | 9-Dec-16 | B |

*, 'II_16N_DICL_top' means that this biofilm sample was collected near station 16, from the top layer of the pipe wall (when biofilm was noticed to be thick), and that pipe is made of ductile iron with cement lining. ^, 'DIPL' refers to ductile iron pipe with polyurethane lining. All the biofilm samples were arranged in the order of the conductivity value.

Table S3 Metadata for the young sensor biofilm samples

| ID | Stn | Date of sampling | ORP (mV) | pH (-) | Conductivity (µS/cm) | Turbidity (FNU) | Water age (hours) | Sample age (days) | Pressure (m) | Pipe diam (mm) | $Cl^-$ (mg/L) | $SO_4^{2-}$ (mg/L) | TOC (mg/L) | TC (mg/L) | TIC (mg/L) | MCA^ (mg/L) | $NH_3$-N (mg/L) | $NO_2$-N (mg/L) | $NO_3$-N (mg/L) | TN (mg/L) | PO4-P (mg/L) |
|---|---|---|---|---|---|---|---|---|---|---|---|---|---|---|---|---|---|---|---|---|---|
| 1 | A16o* | 15-Jun-17 | 337 | 7.39 | 159.11 | 0.76 | 81.45 | 35 | 42 | 300 | 12.42 | 15.71 | 6.14 | 10.43 | 4.29 | 0.24 | 0 | 0.11 | 1.03 | 1.03 | 0.38 |
| 2 | A16 | 1-Nov-17 | 324 | 7.77 | 231.33 | 2.19 | 81.45 | 139 | 41 | 300 | 12.34 | 15.18 | 2.36 | 6.17 | 3.81 | 0.47 | 0 | 0.09 | 1.15 | 1.31 | 0.13 |
| 3 | A14o* | 15-Jun-17 | 446 | 6.65 | 158 | 15[&] | 80.48 | 65 | 61 | 300 | 11.91 | 15.88 | 1.83 | 5.58 | 3.75 | 0.38 | 0 | 0.05 | 1.06 | 1.27 | 0 |
| 4 | A14 | 31-Oct-17 | 212 | 7.3 | 233.62 | 1.4 | 80.48 | 138 | 65 | 300 | 12.52 | 14.95 | 3.52 | 6.82 | 3.3 | 0.5 | 0 | 0.03 | 1.14 | 1.34 | 0.04 |
| 5 | A17 | 22-Nov-17 | 445 | 7.87 | 164.39 | 1.14 | 91.69 | 202 | 60 | 300 | 12.92 | 14.72 | 1.79 | 5.65 | 3.86 | 0.28 | 0 | 0.05 | 1.27 | 1.27 | 0 |
| 6 | B12 | 6-Mar-18 | 329 | 8.98 | 161.99 | 0.23 | 25 | 405 | 31 | 250 | 40.81 | 52.57 | 2.55 | 12.41 | 9.86 | 2.73 | 0.22 | 0 | 0 | 0.46 | 0.02 |
| 7 | B17 | 27-Dec-20 | 390 | 7.88 | 307.61 | 0.19 | 25 | 326 | 26 | 200 | 35.72 | 13.24 | 2.73 | 14.87 | 12.14 | 2.27 | 0.20 | 0.05 | 0 | 0.39 | 0.38 |

Note: *, A16o and A14o refers to the first sampling of YSB at station A16 and A14, respectively. ^ refers to monochloramine. &, the high turbidity value was later found out to be improper positioning of the sensor inside the pipe during the installation. Parameters listed in this table after the pipe diameter column were measured in triplicates and manually.

Table S4. Concentration of orthophosphorus in bulk water at stations A16 and B 17 versus other stations on same campus

| Campus | Station | $PO_4$-P concentration (mg/L) | | Time interval (d) | Turbidity sensor baseline changed |
|---|---|---|---|---|---|
| | | Day when DTM increased | Day when DTM decreased | | |
| A | A16 | 0.38 ± 0.10 | 0.13 ± 0.00 | 91 | Yes |
| | Others | 0.02 ± 0.02[A] | 0.02 ± 0.02[A] | - | No |
| B | B17 | 0.38 ± 0.0 | 0.00 ± 0.00 | 76 | Yes |
| | Others | 0.01 ± 0.01[A] | 0.01 ± 0.01[A] | - | No |

[A] Mean value of all other stations in campus network.

References

Roberts, D.W. (2009) Comparison of multidimensional fuzzy set ordination with CCA and DB-RDA. Ecology 90, 2622-2634.

Torondel, B., Ensink, J.H.J., Gundogdu, O., Ijaz, U.Z., Parkhill, J., Abdelahi, F., Nguyen, V.A., Sudgen, S., Gibson, W., Walker, A.W. and Quince, C. (2016) Assessment of the influence of intrinsic environmental and geographical factors on the bacterial ecology of pit latrines. Microb. Biotechnol. 9, 209-223.
